# A chemogenetic ligand-receptor pair for voltage-gated sodium channel subtype-selective inhibition

**DOI:** 10.1101/2025.11.03.686046

**Authors:** Elizabeth R. Park, Nicholas Denomme, Holly S. Hajare, J. Du Bois

## Abstract

Neuronal excitability relies on the tightly regulated expression and discrete subcellular localization of voltage-gated sodium channels (Na_V_s). These large membrane protein complexes control the movement of sodium ions across cell membranes and are responsible for initiating and propagating action potentials. A desire to better understand the role of Na_V_ subtypes in electrical signal conduction and the relationship between channel dysregulation and specific human pathologies (e.g., epilepsy, musculoskeletal disorders, neuropathic pain) motivates the development of high-precision pharmacological reagents to facilitate Na_V_ studies. Investigations of Na_V_ physiology and nerve cell conduction are limited by a lack of available methods with which to modulate acutely and reversibly the function of individual channel subtypes. Moreover, discriminating between Na_V_s expressed in different cell types is not possible even with potent and selective ligands that target specific channel homologues. We have capitalized on both chemical design and protein engineering to advance a chemogenetic tool to inhibit a single Na_V_ isoform. A synthetic derivative of the bis-guanidinium toxin saxitoxin (STX) is paired with two unique outer pore-forming amino acid mutations to achieve ∼100:1 selectivity for the engineered channel over wild-type Na_V_1.1– 1.4, 1.6, and 1.7. The designer ligand is nanomolar potent against the mutant channel and acts within seconds to block sodium ion conduction; washing cells with buffer solution rapidly and completely restores channel function. This technology will empower studies of Na_V_ physiology and have additional applications for manipulating action potential signals given the requisite role of Na_V_s in electrogenesis.

**SIGNIFICANCE:** Voltage-gated sodium channels are an obligatory component of the biochemical machinery that makes possible electrical signaling in cells. Malfunction of these large protein complexes underlies a number of debilitating human disorders including certain forms of epilepsy, cardiac arrhythmia, and neuropathic pain. A desire to better understand how sodium channels initiate, propagate, and integrate electrical signals in healthy and aberrant cells necessitates access to molecular tools that enable precise manipulation of channel function. This work describes the advancement of such technology, applications of which should facilitate discoveries in foundational and translational research.

## INTRODUCTION

The nervous system is a highly complex, interconnected network of specialized cells that rapidly process and transmit information through chemical and electrical signaling. Electrical impulses (action potentials, APs) arise as frequency- and amplitude-modulated spike trains resulting from the choreographed interactions of ion channels.(1) Among the collection of channels responsible for AP signaling, voltage-gated sodium channels (Na_V_s) provide the explosive inward sodium current that drives AP firing and propagation.(1–3) In mammalian cells, sodium influx is carried by nine isoforms of the pore-forming α-subunit, Na_V_1.1–1.9, each a ∼260 kDa protein with four nonidentical repeats, Domains I–IV (DI–DIV), and co-expressed with one or two auxiliary β-subunits.(4) Na_V_ isoforms are further divided into tetrodotoxin (TTX)-sensitive (Na_V_1.1–1.4, Na_V_1.6, Na_V_1.7) and TTX-resistant (Na_V_1.5, Na_V_1.8, Na_V_1.9) channels based on whether IC_50_ values for TTX fall below or exceed 1 µM.(5) Transcriptional splicing, editing, and post-translational modifications further diversify the population of Na_V_ proteoforms expressed in the plasma membranes of electrically excitable cells.(6–11)

Na_V_ malfunction is causally linked to a number of human pathologies including arrhythmia, chronic pain, epilepsy, and neurodevelopmental disorders.(12–14) A desire to understand how specific Na_V_ subtypes shape APs and how Na_V_ dysregulation underlies disease etiology motivates the work described herein. Recent advances notwithstanding, developing pharmacological tools that target individual Na_V_ subtypes is challenged by the highly conserved nature of the nine α-subunits and the added structural variability introduced through post-transcriptional and post-translational changes.(6, 7, 15) In addition, cell type-selective differentiation of a single Na_V_ isoform is not possible with molecules that target wild-type (WT) channels. An approach that combines chemistry and genetics, analogous to DREADDs and RASSLs for GPCRs or PSAMs for nAChRs,(16–18) offers an enticing solution—one that promises rapid and reversible control of a specifically encoded Na_V_. We report a universal chemogenetic tool for selective inhibition of TTX-sensitive Na_V_s. TTX-sensitive channels appear in the CNS, in cardiac and skeletal muscle, and in non-electrically excitable cells (e.g., lymphocytes, glia).(19, 20) Accordingly, this technology will enable precise pharmacological interrogation of Na_V_-subtype physiology and, given the obligatory role of Na_V_s in AP signaling, offers a complementary method to optogenetics for focally-defined block of neuronal activity.

The highly conserved locus of amino acids that define the outer pore and selectivity filter in Na_V_s offers an ideal receptor site (Site 1) for chemogenetic ligand design (Fig. 1). Organisms that concentrate Site 1 guanidinium toxins, as exemplified by TTX and saxitoxin (STX), introduce select mutations to the outer pore as a resistance mechanism.(21–24) These structural changes do not result in an evident fitness cost nor do they alter gating. As for ligands that target the pore, STX, which binds rapidly, reversibly, and with low nanomolar affinity to TTX-sensitive channels and is accessible along with derivative forms through *de novo* synthesis, is an optimal lead.(25, 26) The overwhelming majority of small molecules that modulate Na_V_s are state-dependent and display slow association and/or dissociation kinetics.(27–34) For these reasons, our studies have focused on compensatory modifications to Site 1 and STX to generate a chemogenetic ligand-channel pair.

**Figure 1.**
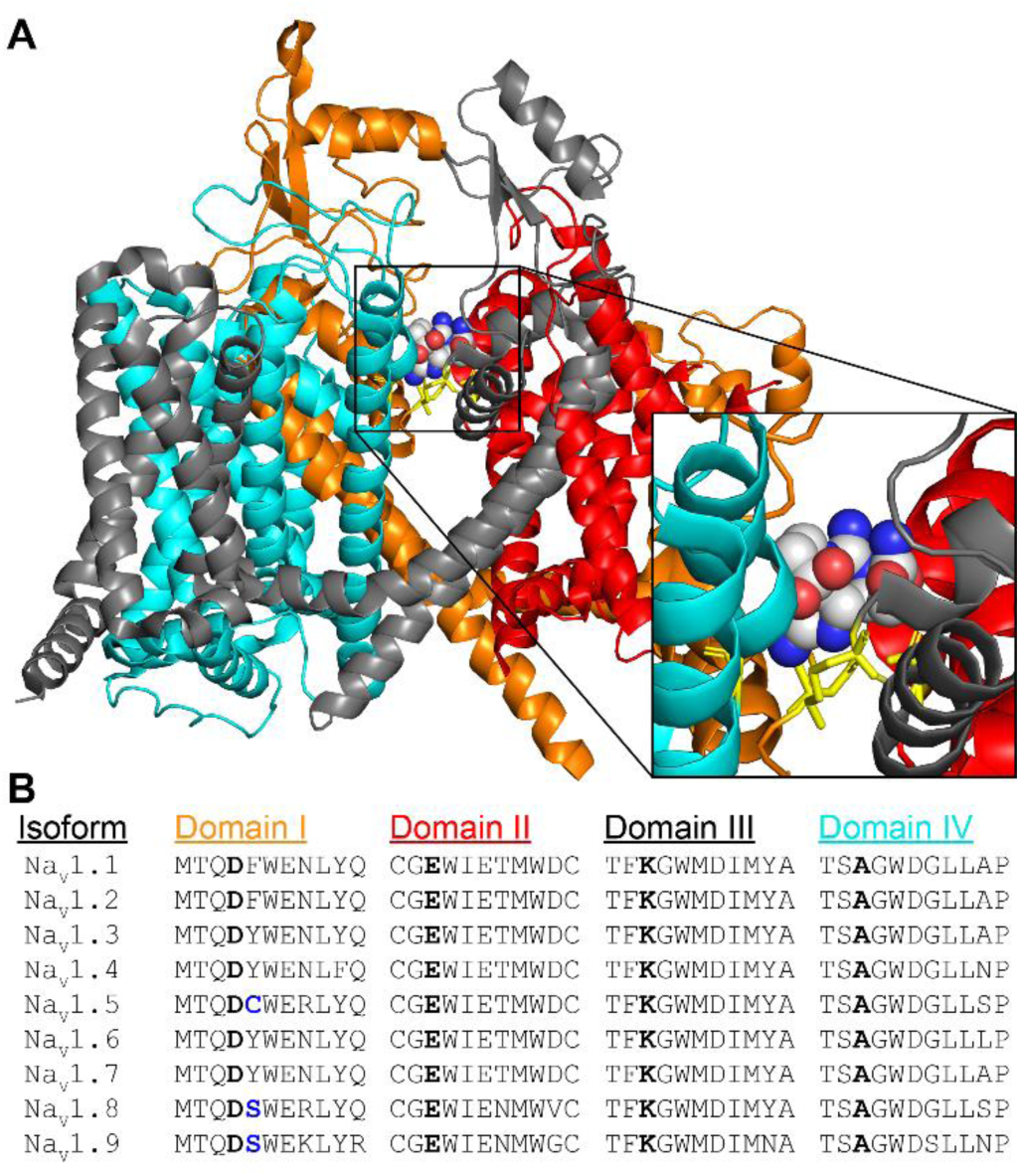
**The structure and sequence of the STX-binding site are highly conserved across voltage-gated sodium channels.** (A) Structure of hNa_V_1.7 with STX bound in the outer vestibule (Site 1). Reproduced from PDB 6J8G.(60) DI = orange, DII = red, DIII = gray, DIV = cyan. The ion selectivity filter is denoted in yellow. (B) Sequence alignment of outer pore residues across rNa_V_1.1–1.9. Ion selectivity filter residues are bolded. Residues that destabilize STX binding to Na_V_1.5, 1.8, and 1.9 are in blue.

## RESULTS

### Evaluation of STX-amides against rNa_V_1.4 as a starting point for chemogenetic tools

For our initial prototype, we targeted a designer molecule that would inhibit an engineered Na_V_ with nanomolar potency (IC_50_ < 100 nM) and >100:1 selectivity relative to WT channels. To identify a starting ligand, we compared STX and STX C13-derivatives that included a naturally occurring acetate ester (LWTX-5), an analogous acetamide **1**, and a small collection of aryl amides.(25, 35) Half-maximal inhibitory concentrations (IC_50_) for both STX and LWTX-5 against WT rat Na_V_1.4 (rNa_V_1.4) α-subunit expressed in Chinese hamster ovary K1 (CHO-K1) cells are 2.9 ± 0.1 and 83 ± 3.9 nM, respectively.(25) A marked decrease in potency, however, is noted across the amide series (Fig. 2*B*), including acetamide **1** (IC_50_ = 1.9 µM). As with STX, channel block by these amide derivatives is fast and reversible. The >500-fold change in IC_50_ values for these compounds is not easily explained by available structural data and may stem from differences in solvation energies and the energetic cost of desolvating the amide moiety. The reason(s) for such potency differences aside, these modified toxins, particularly aryl amides such as STX-NBz **2**, offer an attractive starting point for chemogenetic ligand design. The lack of potency for STX-NBz **2** manifests beyond rNa_V_1.4 and includes all TTX-sensitive channels (Na_V_1.1–1.4, 1.6, 1.7) (Fig. 2*C*).

**Figure 2.**
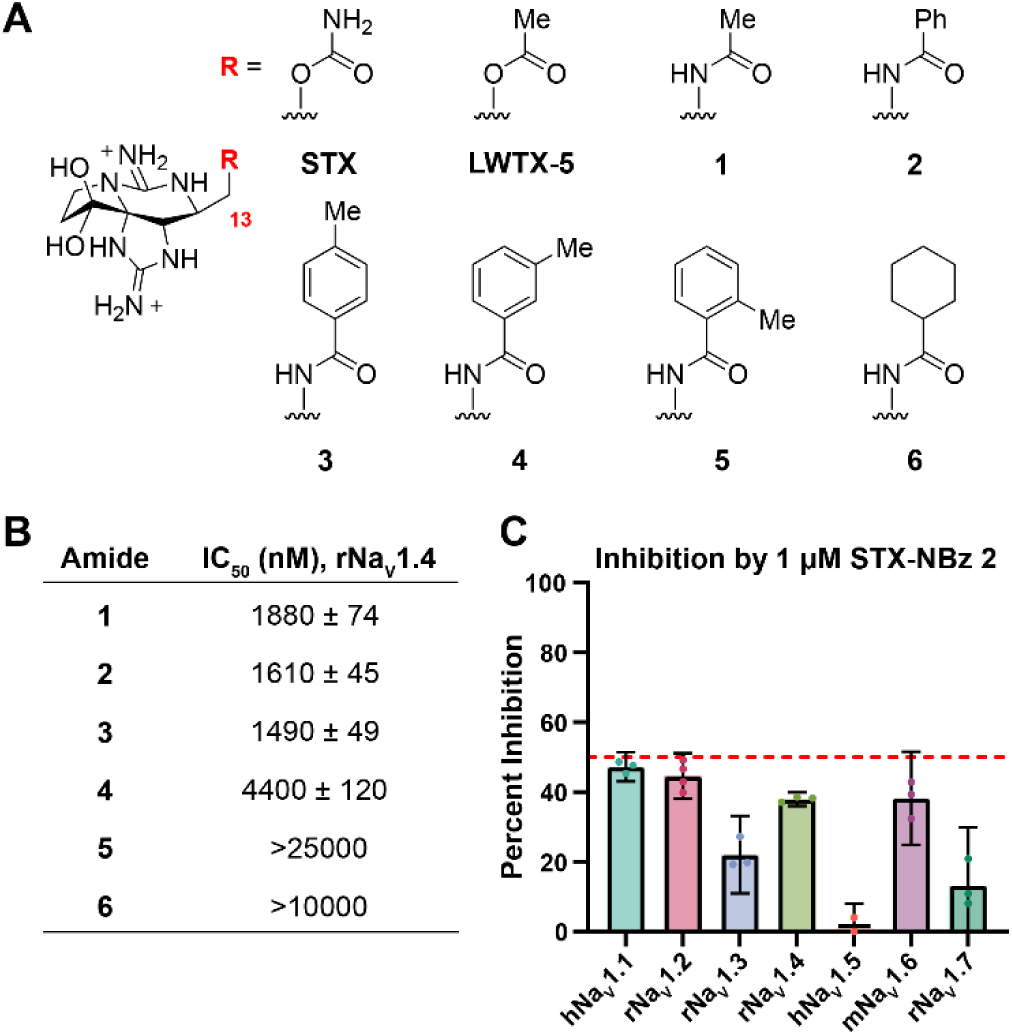
**Electrophysiological characterization of STX-amides against WT Na_V_s.** (A) Structures of STX, LWTX-5, and non-natural STX-amides. (B) Electrophysiological characterization of STX-amides against WT rNa_V_1.4. IC_50_ ± s.e.m. (C) 1 µM STX-NBz **2** was exposed to Na_V_1.1–1.7 to determine cross-isoform potency. Data represent percent reduction in initial current in comparison to baseline, mean ± 95% C.I., n ≥ 3. A red line is drawn at 50% to note IC_50_s are estimated to be at or above 1 µM for all subtypes tested.

### Rescue of Amide Potency through rNa_V_1.4 Outer Pore Mutations

To explore mutations that rescue the potency of STX-amides, single amino acid substitutions were made across DI–DIV in the p-loop α-helices comprising the STX binding site. Because the engineered Na_V_ must retain WT channel function, we opted for conservative mutations in the outer mouth of NaVs, largely targeting residues that vary in species resistant to guanidinium toxins. Mutations were introduced into *Scn4a*, the gene encoding rNa_V_1.4, and the corresponding channels were successfully expressed in CHO-K1 cells. Mutant Na_V_s were tested only if they displayed current densities and gating properties that closely mirrored WT rNa_V_1.4. In the first-pass analysis, the potency of STX-NBz **2** was evaluated against mutant channels at 1 µM (Fig. S3). Known mutations that destabilize STX binding—Y401C, Y401S, E403D, E758D, and D1532N—also disrupted block by STX-NBz **2**, as no recordable inhibition by **2** at 1 µM was measured.(25, 36–38) On the other hand, substitution of D1241 with hydrophobic residues (e.g., Leu, Ile) improved STX-NBz **2** potency (Fig. S3) but decreased the affinity of STX (Fig. S4). Unexpectedly, block of rNa_V_1.4 D762E and G1533A mutants by STX-NBz **2** and STX was enhanced relative to WT channel (Figs. S3, S4); these mutations may similarly influence the binding of other bis-guanidinium toxins.(39) Concentration-response curves with STX-NBz **2** and four rNa_V_1.4 mutant channels, D762E, D1241L, D1241I, and G1533A, show that the potency of STX-NBz **2** is improved 10-fold by the most stabilizing mutation, Domain III D1241L (WT rNa_V_1.4: IC_50_ = 1610 ± 45 nM; D1241L: 163 ± 8 nM, Fig. 3).

**Figure 3.**
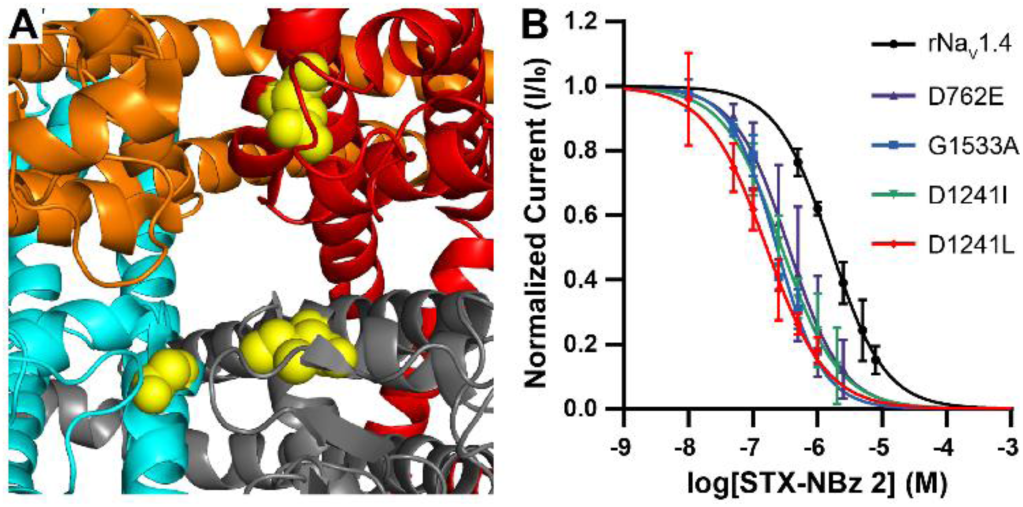
**Mutagenesis of the highly conserved Na_V_ outer pore reveals stabilizing mutations for STX-NBz.** (A) PDB 6J8G (hNa_V_1.7 with STX bound) highlighting the location of STX-NBz **2** stabilizing mutations.(60) DI = orange, DII = red, DIII = gray, DIV = cyan. In sphere model, D762 (DII), I1241 (DIII), and G1533 (DIV) are represented in yellow space filling. (B) Electrophysiological characterization of **2** in transiently expressed rNa_V_1.4 wild-type and select mutant channels. IC_50_s ± s.e.m. (nM), Hill coefficients ± s.e.m.: rNa_V_1.4 = 1610 ± 45, −1.03 ± 0.03; D762E = 357 ± 31, −1.04 ± 0.10; G1533A = 235 ± 11, −1.21 ± 0.07; D1241I = 279 ± 14, −0.97 ± 0.05; D1241L = 163 ± 8, −0.94 ± 0.05. Graph represents mean ± 95% C.I. (rNa_V_1.4, n = 3; D762E, n = 4; G1533A, n = 5; D1241I, n = 3; D1241L, n = 3).

To further boost the potency of STX-NBz **2** as a channel blocker, we investigated Na_V_s in which two stabilizing mutations from different domains were combined in one construct. Of the double mutant channels evaluated, rNa_V_1.4 D762E/G1533A, D762E/D1241L, and D1241L/G1533A were particularly effective in rescuing the potency of the benzamide ligand (Fig. 4*A*). Measured IC_50_ values for STX-NBz **2** against these three mutant channels range between 35–49 nM. We selected the two top performing constructs, rNa_V_1.4 D762E/D1241L and D1241L/G1533A, for subsequent studies. Importantly, introducing either of these pairs of mutations does not alter rNa_V_1.4 functional properties compared to WT, including voltage dependence of activation, inactivation, and ion selectivity (Fig. S5, S6). Inhibition of these double mutant channels by STX-NBz **2** is favored over WT rNa_V_1.4 by ∼45:1.

**Figure 4.**
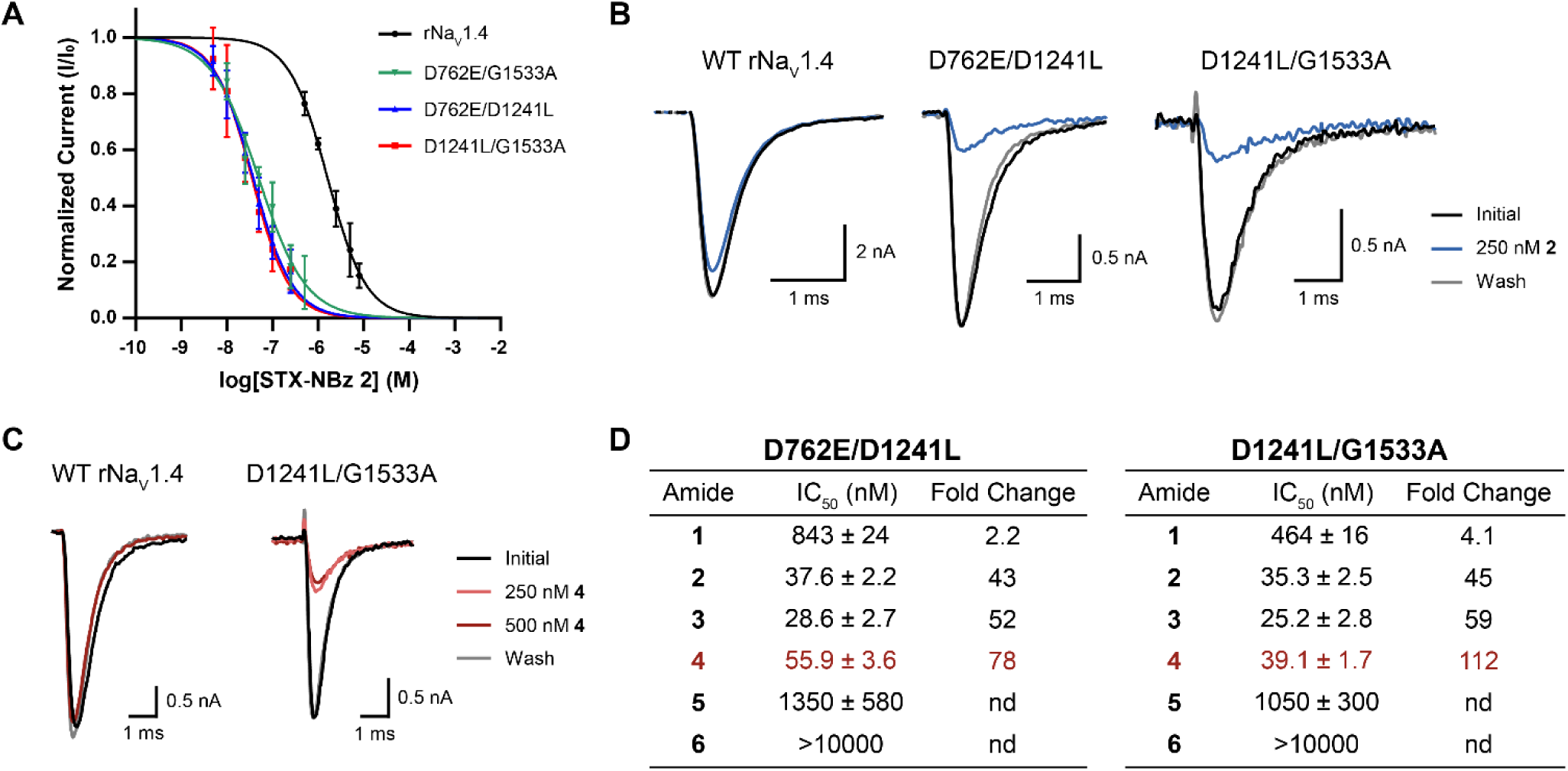
**Two outer pore mutations markedly improve the potency of STX-amides**. (A) Concentration-response curves of STX-NBz **2** in rNa_V_1.4 double mutant channels. IC_50_s ± s.e.m. (nM), Hill coefficients ± s.e.m.: rNa_V_1.4 = 1610 ± 45, −1.03 ± 0.03; D762E/G1533A = 49.3 ± 3.6, −0.84 ± 0.06; D762E/D1241L = 37.6 ± 2.2, −1.01 ± 0.06; D1241L/G1533A = 35.3 ± 2.5, −1.06 ± 0.08. Graph represents mean ± 95% C.I. (rNa_V_1.4, n = 3; D762E/G1533A, n = 3; D762E/D1241L, n = 5; D1241L/G1533A, n = 4). (B) Representative voltage-clamp traces of WT rNa_V_1.4, D762E/D1241L and D1241L/G1533A with 250 nM of **2** applied. Traces collected in the order: Initial, 250 nM, Wash. (C) Representative voltage-clamp traces of WT rNa_V_1.4 and D1241L/G1533A with 250 nM or 500 nM of **4** applied. Traces collected in the order: Initial, 250 nM, 500 nM, Wash. (D) Electrophysiological characterization of all STX-amides against rNa_V_1.4 double mutant channels, D762E/D1241L and D1241L/G1533A, reported as IC_50_ ± s.e.m. (nM). Fold change is reported in comparison to the WT IC_50_ values reported in Figure 2. (**1** vs. D762E/D1241L, n = 3; **1** vs. D1241L/G1533A, n = 4; **3** vs. D762E/D1241L, n = 3; **3** vs. D1241L/G1533A, n = 4; **4** vs. D762E/D1241L, n = 4; **4** vs. D1241L/G1533A, n = 10; **5** vs. D762E/D1241L, n = 4; **5** vs. D1241L/G1533A, n = 3; **6** vs. D762E/D1241L, n = 3; **6** vs. D1241L/G1533A, n = 3). nd = not determined.

### Improving Selectivity through Ligand Modification

We measured IC_50_ values for additional STX-amide derivatives (**1, 3–6**) against rNa_V_1.4 D762E/D1241L and D1241L/G1533A mutants to determine if ligand substitution could bolster the selectivity factor for block versus the WT channel. Among these five ligands, *meta*-toluamide, STX-N^m^Tl **4**, displayed >100:1 selectivity for rNa_V_1.4 D1241L/G1533A over WT and a favorable IC_50_ of 39.1 ± 1.7 nM (Fig. 4*D*). Other C13-amide ligands such as *ortho*-toluamide, STX-N°Tl **5**, and cyclohexylcarboxamide, STX-NCy **6**, were ineffective channel blockers (IC_50_ > 1 µM) or fell short of our targeted 100:1 selectivity (STX-NAc **1**, *para-*toluamide STX-N^p^Tl **3**). The combination of rNa_V_1.4 D1241L/G1533A and STX-N^m^Tl **4** satisfies our original design criteria for a potent and selective chemogenetic tool.

### Evaluating STX-N^m^Tl 4 against alternative Na_V_ subtypes

Given the nearly identical sequence structure of the outer pore-forming p-loop repeats across all Na_V_ subtypes (Fig. 1), we assessed the potency of STX-N^m^Tl **4** against structurally equivalent D/L (DIII)–G/A (DIV) double mutants of Na_V_1.1–1.3 and 1.5–1.7. Sodium current recordings of STX-N^m^Tl **4** with WT Na_V_1.1–1.3 and 1.5–1.7 in CHO-K1 cells confirmed that IC_50_ values for this compound are >2 µM (Fig. 5). As with rNa_V_1.4, introducing D/L–G/A mutations to Na_V_1.1–1.3, 1.6, and 1.7 significantly improves the potency of STX-N^m^Tl **4**, reducing IC_50_ values between 69–149-fold. With the TTX-resistant channel, Na_V_1.5, binding of **4** to Na_V_1.5 D1423L/G1715A is greatly favored over the WT channel, but the potency of this ligand for the double mutant is low (IC_50_ = 2.4 µM). Introducing the D/L-G/A mutants to any one of these subtypes does not alter the biophysical properties of the channel (Fig. S5). Collectively, our electrophysiology data indicate that, in cells expressing both the double mutant and WT Na_V_s, concentrations of STX-N^m^Tl **4** between 250–500 nM should inhibit a large percentage of mutant channels (73–90%) with minimal block of WT conduction (<10%). Like STX, channel block by STX-N^m^Tl **4** is kinetically fast (k_on_ ∼10^6^ M^−1^ s^−1^) and fully reversible (k_off_ ∼10^−2^ s^−1^) (Fig. S2).

**Figure 5.**
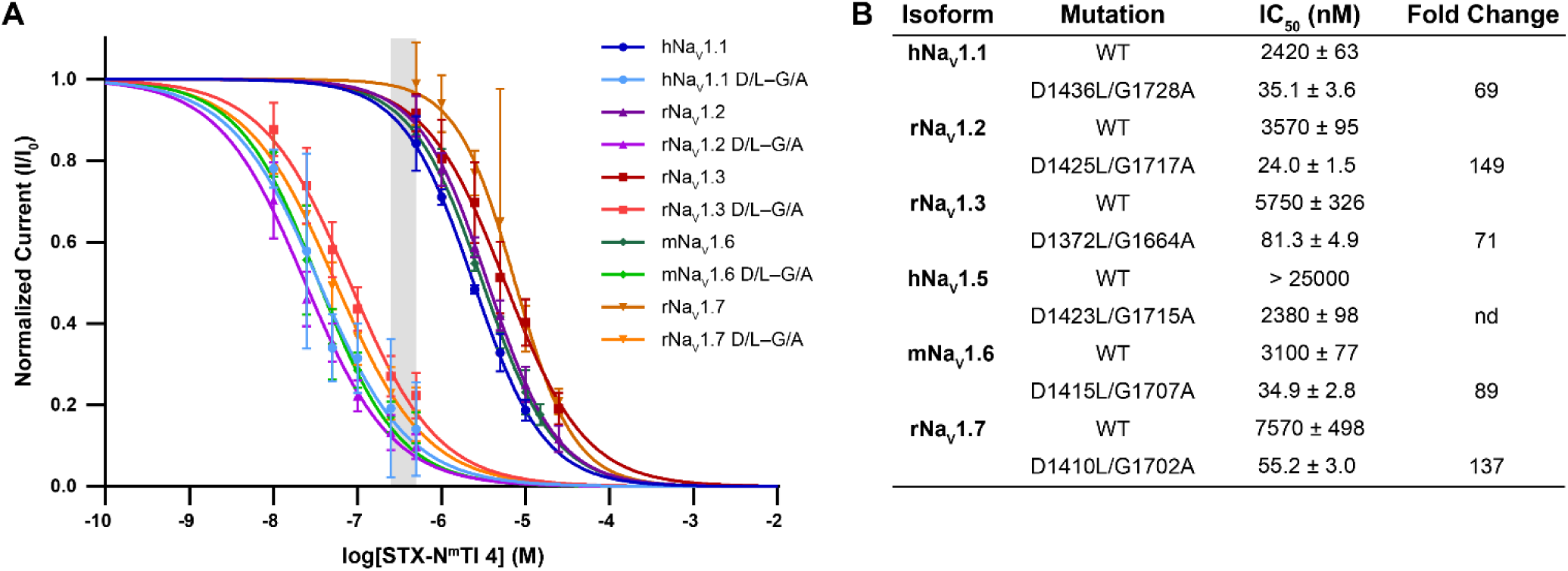
**TTX-‘sensitive’ Na_V_s expressing D/L and G/A outer pore mutations are inhibited selectively by STX-N^m^Tl 4.** (A) Concentration-response curves of **4** with TTX-sensitive WT and double mutant Na_V_1.1–1.7 transiently expressed in CHO-K1 cells. The operative concentration range from 250–500 nM is depicted in gray. Data represents mean ± 95% C.I. (hNa_V_1.1, n = 4; hNa_V_1.1 D/L–G/A, n = 3; rNa_V_1.2, n = 4; rNa_V_1.2 D/L–G/A, n = 6; rNa_V_1.3, n = 4; rNa_V_1.3 D/L–G/A, n = 5; mNa_V_1.6, n = 4; mNa_V_1.6 D/L–G/A, n = 4; rNa_V_1.7, n = 3; rNa_V_1.7 D/L–G/A, n = 5). (B) Table of fold change in IC_50_s comparing WT to double mutant Na_V_1.1–1.7. IC_50_s ± s.e.m.; nd = not determined.

### Effect of STX-N^m^Tl 4 in cultured hippocampal neurons and in slice

To ensure that operating concentrations (250–500 nM) of STX-N^m^Tl **4** do not perturb WT Na_V_ function in primary cells, we applied this ligand to cultured E17–18 rat hippocampal neurons. Sodium current measurements of **4** with hippocampal neurons (days *in vitro*, DIV 6–8) reveals an IC_50_ of 3.72 ± 0.26 µM (Fig. 6*A* and *B*), a value congruent with CHO-K1 cell data and consistent with the expression of multiple TTX-sensitive isoforms (Na_V_1.1–1.3, Na_V_1.6) in these cells.(40–42)

**Figure 6.**
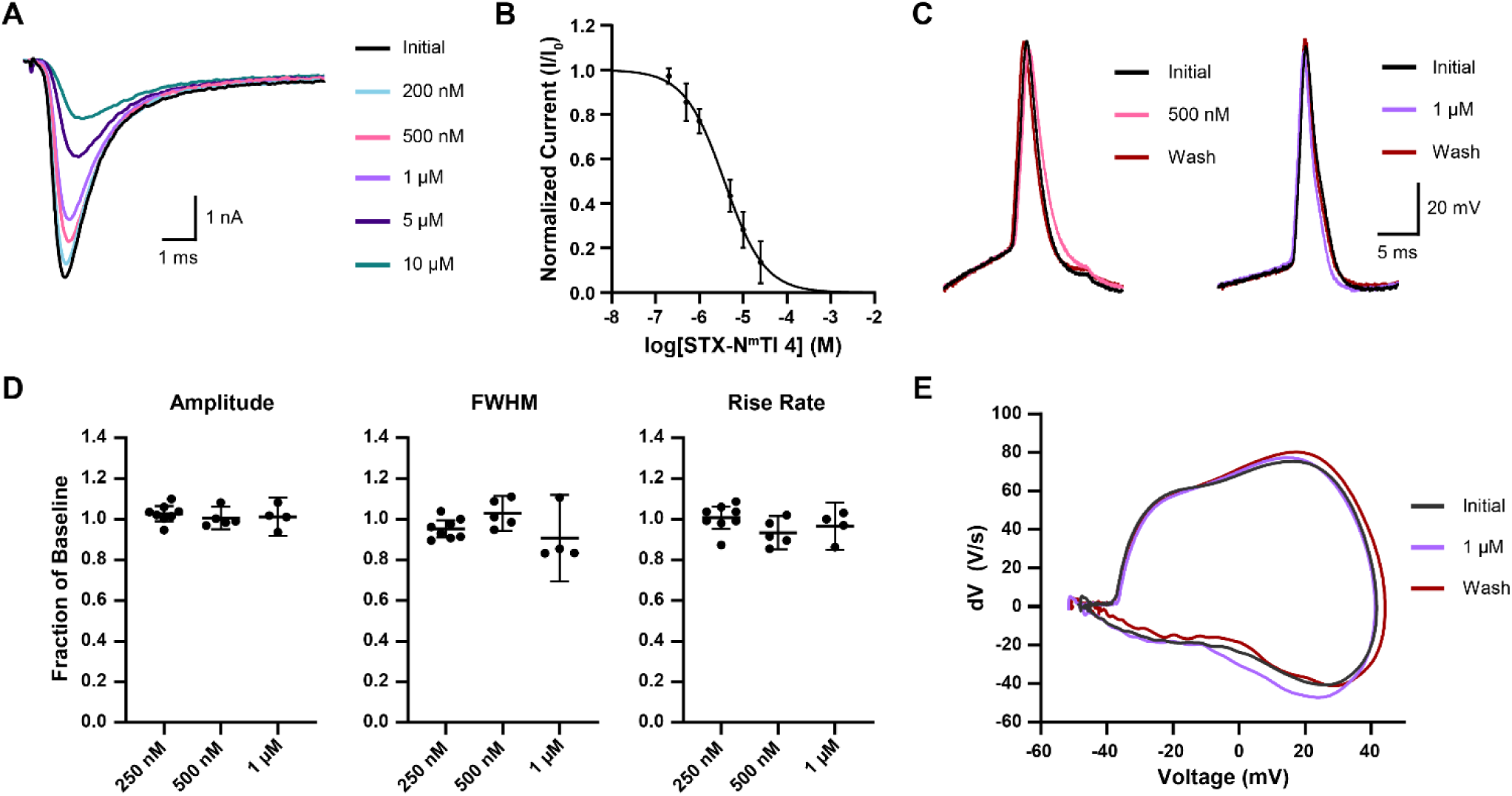
**Effects of STX-N^m^Tl 4 on sodium current and single AP properties in cultured hippocampal neurons.** (A) Representative voltage-clamp traces of rat E17–18 hippocampal cells (DIV 6–8) treated with increasing concentrations of STX-N^m^Tl **4**. Traces collected in the order: Initial, 200 nM, 500 nM, 1 µM, 5 µM, 10 µM. (B) Concentration-response curve of **4** in embryonic hippocampal neurons. IC_50_ ± s.e.m. (nM), Hill coefficient ± s.e.m.: 3720 ± 262, −0.95 ± 0.06. Graph represents mean ± 95% C.I. (n = 7). (C) Representative single AP traces of rat E17–18 hippocampal cells (DIV 9–13) with 500 nM or 1 µM of **4** applied. Traces taken in the order: Initial, 500 nM or 1 µM, Wash. (D) Characteristics of a single elicited action potential with 250 nM, 500 nM, or 1 µM of **4** applied to embryonic hippocampal neurons. Amplitude was recorded as peak mV. Rise rate is recorded as the maximum dV/dt and FWHM represents the full width at half maximum amplitude. Data represent mean ± 95% C.I. Data are not statistically significantly different as determined by paired *t*-tests to baseline values. (For 250 nM, n = 8; 500 nM, n = 5; 1 µM, n = 4). (E) Representative phase plot of a single elicited action potential by an embryonic hippocampal neuron pre-, during, and post-application of 1 µM of **4**. Traces taken in the order: Initial, 1 µM, Wash.

Current-clamp electrophysiology experiments with rat hippocampal neurons (DIV 9–13) were performed using four concentrations of STX-N^m^Tl **4** to determine the influence of this ligand on AP signaling. In these experiments, we evaluated single AP amplitude, half-width, maximum rise rate, and AP train firing rate. Single APs were elicited with a 25 ms injection of +100 pA current. At 1 µM **4**, AP amplitude, half-width, and rise rate were unchanged relative to untreated cells (Fig. 6*C*–*E*) even though at this concentration, ∼30% of WT channels are likely blocked (estimated from voltage-clamp data). This finding is consistent with previous reports that ∼33% of Na_V_s expressed in neurons can be inhibited without altering Na_V_-dependent single AP properties.(43)

Action potential trains were elicited through 750 ms injections of +50–100 pA current. Because repetitive firing requires a reserve of available Na_V_s, the AP firing rate is more sensitive to sodium channel block and >10% inhibition can result in a reduction in firing frequency.(43–46) Application of 250 nM STX-N^m^Tl **4** had no observable influence on AP firing rates (Fig. 7*B* and *C*). Increasing **4** to 500 nM concentration results in an 18% diminution in AP firing rate. Only at 1 µM (∼30% channels blocked) is AP frequency visibly diminished; full ablation of AP trains is observed at 5 µM **4** (∼60% channels blocked). At these concentrations, the effect of **4** on AP firing is completely reversible upon perfusion of cells with external solution.

**Figure 7.**
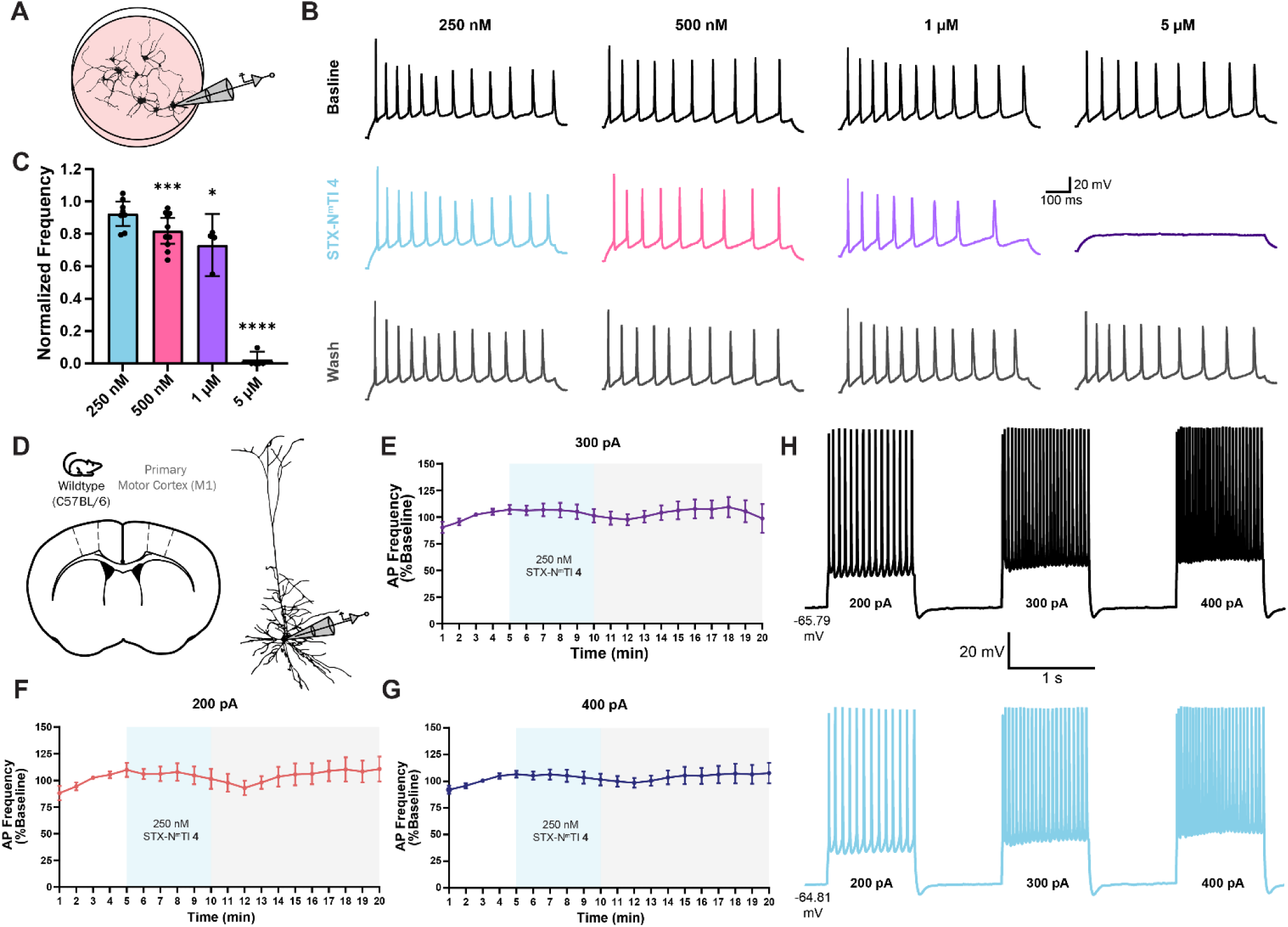
**Effect of STX-N^m^Tl 4 on evoked firing frequency in cultured neurons and cortical slices.** (A) Schematic of a whole-cell recording on cultured embryonic hippocampal cells. (B) Representative current-clamp traces of rat E17–18 hippocampal cells (DIV 9–13) with various concentrations of STX-N^m^Tl **4** applied. Traces collected in the order: Initial, **4**, Wash. Each concentration of **4** was examined on a different cell. (C) Changes in AP firing frequency in an AP train with various concentrations of **4** applied. Data represent mean ± 95% C.I. Statistics were calculated with a paired *t*-test compared to baseline firing frequency, * = p < 0.05, *** = p < 0.001, **** = p < 0.0001. (250 nM, n = 8; 500 nM, n = 10; 1 µM, n = 4; 5 µM, n = 5). (D) Schematic of whole-cell recording configuration in M1 L5 pyramidal neurons. (*E*–*G*) Effects of 250 nM **4** on normalized firing frequency evoked by 200 pA, 300 pA, or 400 pA. Data represents mean ± s.e.m. (N = 3, n = 4). (*H*) Representative AP trains evoked at 200, 300, or 400 pA from the same cell during baseline (black) and 250 nM **4** (blue) conditions.

To further evaluate the working concentration of STX-N^m^Tl **4**, we performed whole-cell current-clamp recordings from layer 5 pyramidal neurons in mouse primary motor cortex (M1). Action potential trains were evoked with somatic current injections (200–400 pA) before, during, and after acute perfusion of **4**. At 250 nM, **4** exhibits no effect on input resistance or evoked firing frequency (Fig. 7*D*–*H*), in keeping with our findings with cultured neurons. Based on our recordings from both hippocampal cultures and cortical slice preparations, 250 nM **4** should serve as a useful operative concentration for both *in vitro* and *ex vivo* electrophysiology experiments, a concentration at which 73–86% of the engineered channels are blocked.

## DISCUSSION

The initiation and propagation of APs require the coordinated action of multiple Na_V_ subtypes. A desire to understand how specific channel isoforms contribute to AP signaling and how AP signaling is altered when subtype trafficking, localization, and/or gating is dysregulated necessitates the invention of high-precision pharmacological tools for altering Na_V_ function. Additionally, the importance of Na_V_s as therapeutic targets to treat peripheral and central nervous system disorders motivates a broader interest in Na_V_ physiology and pathophysiology and inspires the studies herein.(20, 34, 47) In this report, we disclose a chemogenetic mutant Na_V_-ligand pair that offers a general solution for selectively manipulating single Na_V_ subtypes. Future applications of this unique tool are manifold for elucidating Na_V_ subtype function in different cell types and for pharmacologically modulating APs in engineered neuronal cells and tissue.

The advancement of a ‘one-size-fits-all’ designer ligand-Na_V_ mutant combination to interrogate the physiology of individual channel isoforms follows from the availability of bis-guanidinium toxin derivatives through *de novo* synthesis and the highly conserved outer vestibule of Na_V_ subtypes. Saxitoxin is an optimal template for chemogenetic ligand design given its desirable pharmacodynamic properties (i.e., reversible and state-independent channel binding). The amenability of the STX binding site (Site 1) to select mutations that cause little to no evident perturbation to channel function is equally critical to the advancement of this work.

We have identified three amino acids D762 (DI), D1241 (DIII), and G1533 (DIV) (rNa_V_1.4 numbering) that, when appropriately mutated, stabilize binding of STX-N^m^Tl **4** and related amide structures. When evaluated against WT Na_V_1.1–1.4, 1.6, and 1.7 (TTX-sensitive channels), inhibition of double mutant D/L (DIII)–G/A (DIV) constructs by STX-N^m^Tl **4** is 69–149x more selective. In addition to its potency (IC_50_ values between 24–81 nM), binding of **4** to Na_V_s is fast and fully reversible. At 250–500 nM concentrations of **4**, levels that are near saturation block of the D/L-G/A double mutant (73–90%), the influence of this ligand on WT channels in CHO-K1 cells, primary cultured neurons, and mouse brain slice is effectively nil.

Like STX, STX-N^m^Tl **4** is a state-independent channel antagonist. This property is advantageous as it enables block of Na_V_ conductance without stimulation using electrophysiological protocols or pharmacological agents (e.g., peptide venoms) to access different conformational states of the channel. Channel block occurs within seconds upon application of **4**, thereby allowing acute and specific inhibition of mutant Na_V_s in the complex electrical circuit of the neuron. A related approach that enables differentiation of Na_V_1.2 or Na_V_1.6 from other Na_V_ isoforms relies on select mutations in the DIV voltage-sensing domain and a state-dependent arylsulfonamide compound (ASC).(48, 49) Mutant channel block by the ASC, however, is incomplete and requires an intricate voltage protocol to enable prolonged depolarization of the cell. These constraints and the use-dependent activity of such ligands can complicate interpretation of the electrophysiological data. The STX-N^m^Tl–Na_V_ D/L–G/A double mutant pair is advantaged in this regard.

The ability to block sodium conductance resulting from a single Na_V_ isoform in a specific cell type should empower studies of cellular, circuit, and network physiology. Modern genetic methods make possible cell type-specific knock-in of D/L–G/A double mutants for any TTX-sensitive WT Na_V_ channel of interest including those present in glial, cancer, or other non-excitable cells. Differentiating between Na_V_ isoforms expressed in more than one cell type is not possible with inhibitors such as PF-05089771 or VX-548,(33, 50, 51) which target WT channels—only a chemogenetic tool makes such precision possible. While experiments involving genetic knock-out or knock-down of Na_V_s have provided valuable insights into subtype function, compensatory effects (i.e., upregulation of an alternative isoform) can often obscure results.(52–55) The introduction of point mutations at Site 1 to a TTX-sensitive WT Na_V_, however, should not perturb channel expression and cellular function.

As TTX-sensitive Na_V_s are ubiquitously expressed in CNS tissue and obligatory for AP signaling, fast and reversible pharmacological channel block will enable frequency and amplitude modulation of electrical signals in genetically-defined neurons that comprise a specific neural circuit. Opportunities to interrogate the role of Na_V_ subtypes in glia and how selective modulation of these channels alter APs are also possible. Ultimately, we expect chemogenetic, STX-based tools to facilitate efforts to understand how Na_V_ subtypes orchestrate electrical signaling and signal integration in development and disease.

## MATERIALS AND METHODS

### Synthesis

The synthesis of saxitoxin derivatives was previously described.(25, 39, 44) For characterization of STX-amides, see *SI Appendix, Materials and Methods*.

### Plasmids

A human Na_V_1.1 (hNa_V_1.1) construct, pIR-CMV-SCN1A-Variant-1-IRES-mScarlet, was a gift from Dr. Al George (Northwestern University Feinberg School of Medicine, Department of Pharmacology) (Addgene plasmid #162278 ; http://n2t.net/addgene:162278 ; RRID:Addgene_162278).(56) CMV-promoted FLAG-rNa_V_1.2, rNa_V_1.3, and myc-mouse Na_V_1.6 (myc-mNa_V_1.6) in modified pcDNA3.1(+) vectors were constructed by VectorBuilder (Chicago, IL) or GenScript (Piscataway, NJ). The full-length cDNA used for excision and insertion into low-copy modified pcDNA3.1(+) were from oocyte expression vectors provided by Dr. A. L. Goldin (University of California, Irvine, Department of Microbiology & Molecular Genetics).(44, 57) MT1-promoted rNa_V_1.4 in a pZem288 vector was provided by Dr. S. Rock Levinson (University of Colorado, School of Medicine, Department of Physiology and Biophysics).(58) CMV-promoted hNa_V_1.5 in the mammalian expression vector pcDNA3.1(+) was gifted by Dr. T. R. Cummins (Indiana University, Department of Biology). CMV-promoted rNa_V_1.7 (Accession No. NM_133289.1) in a pcDNA3.1(+) vector was purchased from GenScript. All plasmids were amplified in DH5α (Thermo Fisher, Waltham, MA) or stbl2 (Thermo Fisher) competent cells grown at 37 °C and purified using the QIAprep Spin Miniprep Kit (Qiagen, Germantown, MD) according to the manufacturer’s instructions. Due to the inherent instability of the plasmid, the rNa_V_1.7 wild-type construct and mutant channels were prepared in CopyCutter EPI400 Competent Cells (Biosearch Technologies, Hoddesdon, United Kingdom) at 30 °C.

### Mutagenesis

Na_V_ mutants were prepared by site-directed mutagenesis using the QuikChange Lightning Site-Directed Mutagenesis Kit (Agilent Technologies, Santa Clara, CA) according to the manufacturer’s instructions. Primers (SI Appendix, Materials and Methods) were synthesized by Elim Biopharmaceuticals, Inc. (Hayward, CA). Product plasmids were amplified and purified as described above. Following purification, mutations were confirmed through DNA sequencing with Elim Biopharmaceuticals, Inc. or Plasmidsaurus (South San Francisco, CA). Double mutant channels rNa_V_1.3 D1372L/G1664A and rNa_V_1.7 D1410L/G1702A were prepared by GenScript and amplified and purified as described previously.

### Chinese Hamster Ovary (CHO) Cells

Chinese hamster ovary-K1 (CHO-K1) cells (CCL-61TM) were purchased from American Type Culture Collection (ATCC, Manassas, VA). Cells were grown on 10-cm tissue culture dishes in F12-K medium (ATCC) supplemented with 10% fetal bovine serum (ATCC) and 50 U/mL penicillin-streptomycin (Thermo Fisher). Cells were maintained at 37 °C with 5% CO_2_ and passaged according to previously described protocols.(44)

For electrophysiology, CHO-K1 cells were grown in 6- or 24-well tissue culture dishes and generally transfected at 40–80% confluency. Cells were co-transfected with Na_V_ plasmid (6-well: 2–4 μg; 24-well: 500–700 ng) and enhanced green fluorescent protein (GFP) plasmid (6-well: 0.8–1.2 μg; 24-well: 80–100 ng) using Lipofectamine^TM^ LTX with PLUS^TM^ Reagent (Thermo Fisher) according to the manufacturer’s instructions. GFP was omitted from the transfection of hNa_V_1.1 as the construct contains IRES-mScarlet. Following transfection, cells were maintained at 37 °C with 5% CO_2_ for 14–48 hours. Transfected cells were washed with PBS, dissociated using trypsin-EDTA, diluted with complete media, and then plated onto 5 mm No. 1 cover glass slips (Warner Instruments, Holliston, MA) 1–2 hours prior to whole-cell patch clamp electrophysiology

### Rat embryonic day 18 Sprague Dawley hippocampal neurons

Prior to dissection, 5 mm diameter, 0.15 mm-thick round glass coverslips (Warner Instruments) were coated overnight with 1 mg/mL poly-D-lysine hydrobromide (PDL, molecular weight 70,000–150,000 Da, Millipore Sigma) in 0.1 M, pH 8.5 borate buffer in a 37 °C, 5% carbon dioxide, 96% relative humidity incubator.

Hippocampi were dissected from embryonic day 17–18 fetuses into ice-cold Hibernate E (BrainBits, LLC, Springfield, IL) as previously described.(59) Following dissection, cells were dissociated in 2 mL of trypsin-EDTA for 15 min at 37 °C. Subsequently, trypsinized cells were quenched with 10 mL of quenching medium (DMEM high glucose (Thermo Fisher) supplemented with 15% fetal bovine serum, 100 U/mL penicillin–streptomycin, and 1 mM MEM sodium pyruvate (Atlanta Biologicals, Flowery Branch, GA)). The tissue was allowed to settle, the supernatant was removed, and the tissue pellet was rinsed three times with 5 mL of quenching medium. Cells were then manually triturated into 2 mL of plating medium (DMEM supplemented with 10% FBS, 50 U/mL penicillin–streptomycin, and 1 mM MEM sodium pyruvate) by pipetting with a fire-polished 9” borosilicate glass Pasteur pipet (Thermo Fisher).

Cells were plated onto PDL-coated 5 mm glass coverslips in 35 mm tissue culture dishes containing 2 mL of plating medium at a density of 200,000 cells/dish (for voltage-clamp experiments) or 500,000 cells/dish (for current-clamp experiments). After 45 min, coverslips were transferred to new tissue culture dishes containing 2 mL of maintenance medium (neurobasal supplemented with 1x B-27 Supplement, 1x Glutamax, and 50 U/mL penicillin–streptomycin (Thermo Fisher)). Cells were fed every 3–4 days by replacing 2/3 of the conditioned medium.

### Sodium current recordings

All voltage-clamp recordings were acquired at 100 kHz with an Axopatch 200B amplifier and digitized with a Digidata 1322A (Molecular Devices, San Jose, CA). Signals were filtered with a built-in lowpass, four-pole Bessel filter with a cutoff of 5 kHz. For CHO-K1 cells, fire-polished, thin-walled borosilicate pipettes (1.0–5.0 MΩ) contained an internal solution composed of 40 mM NaF, 1 mM EDTA, 20 mM HEPES, and 125 mM CsCl (pH 7.4 with 50 wt. % aqueous CsOH, 330–335 mOsm). The external solution comprised 160 mM NaCl, 2 mM CaCl_2_, and 20 mM HEPES (pH 7.4 with 50 wt. % aqueous CsOH, 325– 330 mOsm).

Voltage-clamp recordings in cultured hippocampal neurons were performed at DIV 6–8 using fire-polished, thin-walled borosilicate pipettes (3–10 MΩ) with an internal solution composed of 114.5 mM gluconic acid, 114.5 mM CsOH, 2 mM NaCl, 8 mM CsCl, 10 mM MOPS, 4 mM EGTA, 4 mM MgATP, and 0.3 mM Na_2_GTP (pH 7.3 with CsOH, 260–265 mOsm). The external solution was Hibernate E low fluorescence (BrainBits, Springfield, IL) (260–275 mOsm with NaCl).

The membrane was kept at a holding potential of –100 mV for CHO-K1 cells and –80 mV for neurons unless otherwise noted. Prior to recording, cells were typically equilibrated for >5 min after establishing whole-cell configuration. Series resistance was compensated at 80–95% with a τ_lag_ of 20–30 or 35 ms for CHO-K1 cells or hippocampal neurons, respectively. Leak currents were subtracted using a standard P/4 protocol of opposite polarity. All measurements were recorded at room temperature (20–25 °C). Data were collected on cells exhibiting >0.75 nA current upon depolarization (unless otherwise noted), leak current <0.2 nA, and I/I_0_ <15% of I_0_ upon STX or STX-amide wash-out. Due to low current density in both WT and mutant rNa_V_1.3 channels, data were collected on cells exhibiting >0.5 nA of current. Specific protocols for each experiment are given below or in *SI Appendix, Materials and Methods*.

### Screening and Concentration-response Curves

Currents were elicited by 10 ms step depolarizations from a holding potential (–100 mV for CHO-K1 cells expressing hNa_V_1.1, rNa_V_1.2, rNa_V_1.3, rNa_V_1.4, mNa_V_1.6, or rNa_V_1.7; –120 mV for CHO-K1 cells expressing hNa_V_1.5; or –80 mV for E18 hippocampal neurons) to 0 mV at a rate of 0.5 Hz. Toxin solutions were perfused into the bath at a rate of 1 mL/min in the order of lowest to highest concentration through manual perfusion. Cells were bathed in each toxin concentration for 60 seconds with continuous recording to allow complete wash on and equilibration. External solution was applied as a control after the first three concentrations and after all concentrations were applied to observe complete ligand wash-out within 15% of initial peak current.

Data were normalized to control currents (I_0_), plotted against toxin concentration, and analyzed using Prism 10.5 (GraphPad Software, LLC, San Diego, CA). For screening results, percent reduction in initial current is plotted. For concentration-response curves, data were fit to a dose-response curve with variable slope and bottom and top parameters constrained to 0 and 1 respectively. The number of observations (*n*) was ≥3 for all reported data. Errors in IC_50_ values and Hill coefficient values are expressed as s.e.m.

### Action potential recording

#### Cultured hippocampal neurons (DIV 9–13)

Current clamp recordings were acquired at 100 kHz and filtered with a built-in lowpass, four-pole Bessel filter with a cutoff of 10 KHz. The internal solution was composed of 105 mM potassium gluconate, 30 mM KCl, 10 mM HEPES, 10 mM Na_2_phosphocreatine, 1 mM EGTA, 4 mM MgATP, and 0.3 mM Na_2_GTP (pH 7.4 with KOH, 275–280 mOsm); the external solution comprised Hibernate E low fluorescence adjusted to 285–300 mOsm with NaCl. Cells with resting membrane potentials of −50 to −70 mV, not varying more than ±5 mV over the course of the experiment, with 0 to −100 pA of injected holding current were used for experiments. Series resistance was typically compensated at 90–95% with a τ_lag_ of 35 ms.

### Single AP Recordings

A single action potential was elicited by a 25 ms current injection to +100 pA at a rate of 0.2 Hz. A 200 ms injection of −30 pA was included after the elicited AP to test input resistance. Five traces were recorded at baseline, with STX-N^m^Tl **4** applied, and following wash-out of **4**. Amplitude, full width at half maximum, and maximum dV/dt were calculated in Clampfit for each trace, and values were normalized to the average baseline values. Cells were excluded if, after wash-out, both amplitude and full width at half maximum exceeded ±15% of the baseline value.

### AP Trains

Action potentials were elicited by a 750 ms current injection of +50–100 pA at a rate of 0.125 Hz. A 500 ms injection to −30 pA was included prior to the elicited AP trains to test input resistance. Cells were selected for analysis based on firing a minimum of six APs per 750 ms. The number of APs were counted in five traces before application of STX-N^m^Tl **4**, after application, and post wash-out. The average number of APs recorded under each condition were normalized to the average number of APs in the baseline. Cells were excluded if, after wash-out, the number of APs fired exceeded ±15% of the baseline value.

#### Acute brain slices

Acute brain slices were prepared from adult male and female C57BL/6 mice between postnatal day P14–21. Mice were anesthetized with isoflurane anesthesia and transcardially perfused prior to decapitation. Brains were carefully extracted and 250 µm coronal slices were cut using a vibratome (Leica VT1200 S) in ice-cold slice solution containing (in mM): 110 sucrose, 62.5 NaCl, 2.5 KCl, 6 MgCl_2_, 1.25 KH_2_PO_4_, 26 NaHCO_3_, 0.5 CaCl_2_, 20 D-glucose, pH 7.35–7.40. Slices were incubated in slice solution for 20 min at 35 °C then transferred to a recovery chamber with artificial cerebrospinal fluid (ACSF) containing (in mM): 125 NaCl, 2.5 KCl, 1 MgCl_2_, 1.25 KH_2_PO_4_, 26 NaHCO_3_, 2 CaCl_2_, and 20 D-glucose, pH 7.35–7.40.

Brain slices were stored in room temperature ACSF and used for recording 1–5 hours later. All solutions were saturated with carbogen. Individual brain slices were placed in an RC-27 recording chamber (Warner Instruments) and superfused with carbogen-saturated ACSF (33–36 °C) at a flowrate of 2–3 mL/min. Layer 5 pyramidal neurons in the primary motor cortex (M1) were visualized with a BX51WI upright microscope (Olympus, Center Valley, PA) equipped with IR-DIC optics and a 40X 0.8 NA water immersion objective (LUMPLFLN, Olympus). Whole-cell recordings were made using borosilicate pipettes (4–7 MΩ) filled with internal solution containing (in mM): 130 K-gluconate, 10 KCl, 10 Na-phosphocreatine, 4 Mg-ATP, 0.3 Na-GTP, 10 HEPES, 290 mOsm, pH 7.2 adjusted with KOH. STX-N^m^Tl **4** was dissolved in water and used for recording within 24 hours. To determine the acute effects of STX-N^m^Tl **4** on the somatic intrinsic excitability in M1 layer 5 pyramidal neurons, continuous AP trains were evoked using 1s current injections ranging from 100–400 pA during 5 min of baseline (ACSF), during 10 min of STX-N^m^Tl **4** perfusion, and 10 min of wash-out (ACSF). Resting membrane potential at the start of the recordings was between −60 and −80 mV. A small amount of holding current (±150 pA) was injected to bring each cell close to −70 mV prior to AP recordings. Input resistance was monitored throughout the duration of all recordings using a −50 pA hyperpolarizing pulse. Recordings were not included in the final analysis if input resistance changed by >20%. Pipette capacitance neutralization and bridge balance were manually adjusted prior to current-clamp recordings. The liquid junction potential was not corrected. Current-clamp recordings in slice were sampled at 20 kHz and filtered at 10 kHz using a MultiClamp 700B and Digidata 1550B and analyzed using pClamp11 (Molecular Devices). Data are presented as mean value ± s.e.m., where n or N represent number of cells or animals, respectively.

### Data Analysis

All data analysis was performed using the software packages Clampfit v10.5 (Molecular Devices), Microsoft Excel, or Prism v10.5. The statistical significance of differences between mean values was evaluated using Student’s *t*-test, paired *t*-test, or one-way ANOVA with Dunnett’s correction with p < 0.05 considered significant. Results are presented as mean ± 95% C.I. unless indicated otherwise. P-values are denoted as ns (not significant), p > 0.05; *, p ≤ 0.05; **, p ≤ 0.01; ***, p ≤ 0.001; and ****, p ≤ 0.0001.

### Ethics oversight

All animal care and dissection procedures were approved by the Stanford Institutional Animal Care & Use Committee (IACUC).

## Acknowledgements

The authors thank Roshni Bhaskar for her assistance with initial studies to prepare STX amide ligands, Dr. Erica Liu and Prof. Bianxiao Cui for help with neuron preparations, and Dr. Torben Neelands and Steven Miller for advice with electrophysiology experiments. This work was supported by National Institutes of Health (NIH) R01-GM117263. E.R.P. was supported by the Stanford ChEM-H Chemistry/Biology Interface Program, the National Institute of General Medical Sciences (NIGMS) of the NIH under Award Number T32-GM120007, and the National Sciences Foundation (NSF) Graduate Research Fellowship Program. H.S.H. was supported by the Department of Defense (DOD) National Defense Science and Engineering Graduate (NDSEG) Fellowship. This work utilized the Thermo Exploris 240 LC/MS system (RRID:SCR_022216) that was purchased with funding from Stanford C-ShaRP (RRID:SCR_022986) and was supported by Stanford University Mass Spectrometry (RRID:SCR_017801).

## Data Availability

The datasets generated and analyzed in this study are available from the corresponding author upon reasonable request.

## Author Contributions

E.R.P, H.S.H, and J.D. conceived of the project. E.R.P, N.D., and H.S.H conducted experiments. E.R.P, N.D., and J.D. were involved in analyzing the data. E.R.P and J.D wrote the paper with input from N.D. and H.S.H.

