## Supplemental Information for "A chemogenetic ligand-receptor pair for voltage-gated sodium channel subtype-selective inhibition"

**This PDF file includes:**

Figures S1 to S7

SI Appendix, Materials and Methods

SI References

### Figures

**Figure S1.** Evaluating the potency of STX-amides **1**, **3**–**6** against WT rNa$_V$1.4 and rNa$_V$1.4 D762E/D1241L and D1241L/G1533A double mutant channels.

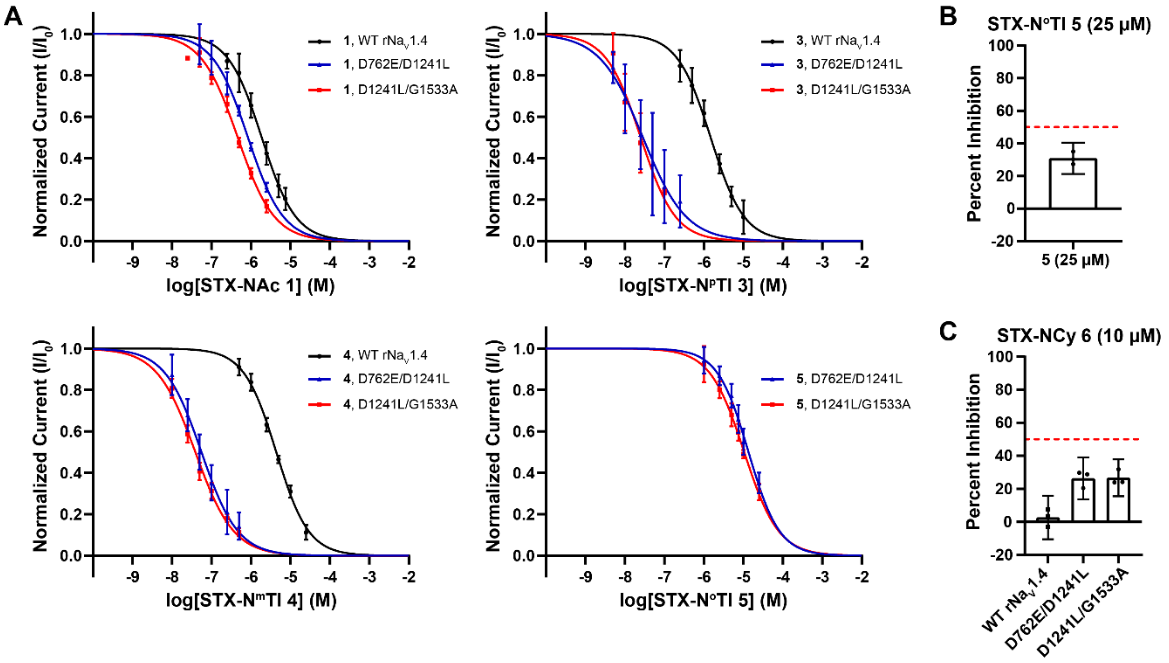


(*A*) Concentration-response curves of STX-amides (**1**, **3**–**5**) against WT and double mutant rNa$_V$1.4. $IC_{50}$ values for WT are reported in Figure 2B; $IC_{50}$ values for double mutant channels are reported in Figure 4D. Hill coefficients ± s.e.m.: **1** WT, −1.01 ± 0.04; **1** D762E/D1241L, −0.97 ± 0.03; **1** D1241L/G1533A, −0.94 ± 0.03; **3** WT, −1.03 ± 0.03; **3** D762E/D1241L, −0.80 ± 0.07; **3** D1241L/G1533A, −0.99 ± 0.12; **4** WT, −1.06 ± 0.03; **4** D762E/D1241L, −0.94 ± 0.06; **4** D1241L/G1533A, −0.90 ± 0.04; **5** D762E/D1241L, −1.11 ± 0.06; **5** D1241L/G1533A, −1.00 ± 0.04. Graph represents mean ± 95% C.I. (**1** WT, n = 4; **1** D762E/D1241L, n = 3; **1** D1241L/G1533A, n = 4; **3** WT, n = 3; **3** D762E/D1241L, n = 3; **3** D1241L/G1533A, n = 4; **4** WT, n = 4; **4** D762E/D1241L, n = 4; **4** D1241L/G1533A, n = 10; **5** D762E/D1241L, n = 4; **5** D1241L/G1533A, n = 3).

(*B*) A concentration-response curve for STX-N$^o$Tl **5** was not obtained given the poor potency of this ligand. Block of WT rNa$_V$1.4 by compound **5** was evaluated at a single concentration (25 mM). The dashed red line represents 50% channel block. Data represents mean ± 95% C.I. (n = 3).

(*C*) A concentration-response curve for STX-NCy **6** was not obtained given the poor potency of this ligand. Block of WT rNa$_V$1.4 and double mutant channels, D762E/D1241L and D1241L/G1533A, by compound **6** was evaluated at a single concentration (10 mM). The dashed red line denotes 50% channel block. Data represents mean ± 95% C.I. (for WT rNa$_V$1.4, n = 3; rNa$_V$1.4 D762E/D1241L, n = 3; rNa$_V$1.4 D1241L/G1533A, n = 3).

**Figure S2.** Kinetics of inhibition for STX-N$^{m}$Tl **4** against rNa$_{V}$1.4.

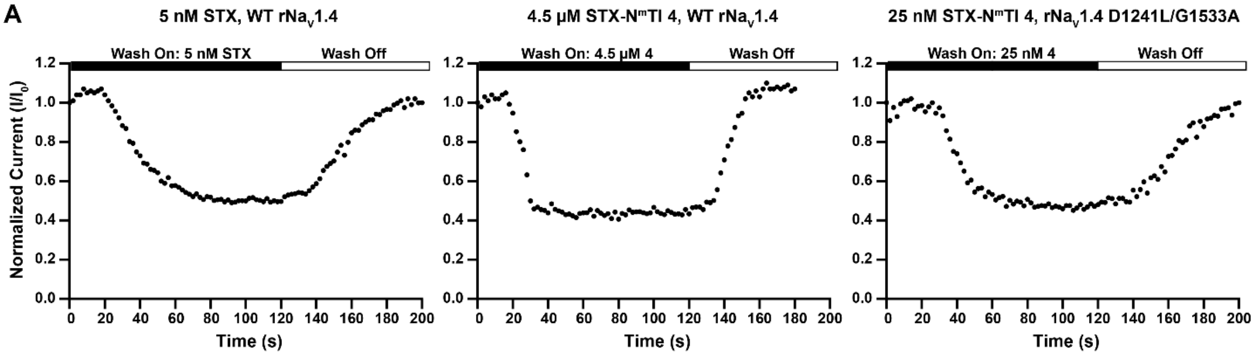


B

| Compound (Concentration) | $I/I_0$ | $k_{on}$ $M^{-1}s^{-1}$ | $k_{off}$ $s^{-1}$ |
|---|---|---|---|
| STX (5 nM) | 0.51 | $2.9 \pm 0.4 \times 10^{6}$ | $2.2 \pm 0.1 \times 10^{-2}$ |
| **4** (4.5 μM) | 0.46 | $2.0 \pm 0.4 \times 10^{4}$ | $2.1 \pm 0.6 \times 10^{-2}$ |
| **4** (25 nM, D/L–G/A) | 0.56 | $1.8 \pm 0.2 \times 10^{6}$ | $7.4 \pm 0.3 \times 10^{-3}$ |

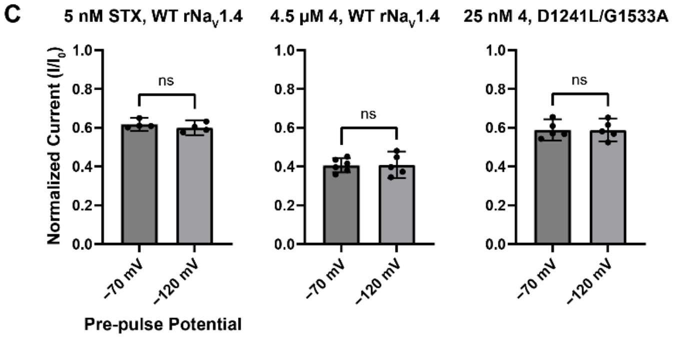


(*A*) Representative current vs time graphs of wash on/off experiments for STX and STX-N$^{m}$Tl **4**. Both compounds exhibit complete wash on within seconds of bath exchange.

(*B*) Table of calculated $k_{on}$ and $k_{off}$ for STX and **4**. Data represent mean ± s.e.m.; for STX, n = 4; **4** against WT rNa$_{V}$1.4, n = 4; **4** against rNa$_{V}$1.4 D1241L/G1533A, n = 3.

(*C*) State-dependence of binding of STX and **4** against WT rNa$_{V}$1.4. CHO-K1 cells transiently expressing rNa$_{V}$1.4 were held at −70 mV or −120 mV for 200 ms prior to depolarization to 0 mV to allow channels to sample different conformational states. Data represent mean ± 95% C.I., n ≥ 4. Statistics were calculated with a two-tailed student *t*-test, ns = p value > 0.05.

**Figure S3.** Screening STX-NBz **2** against rNa$_V$1.4 DI–IV mutations.

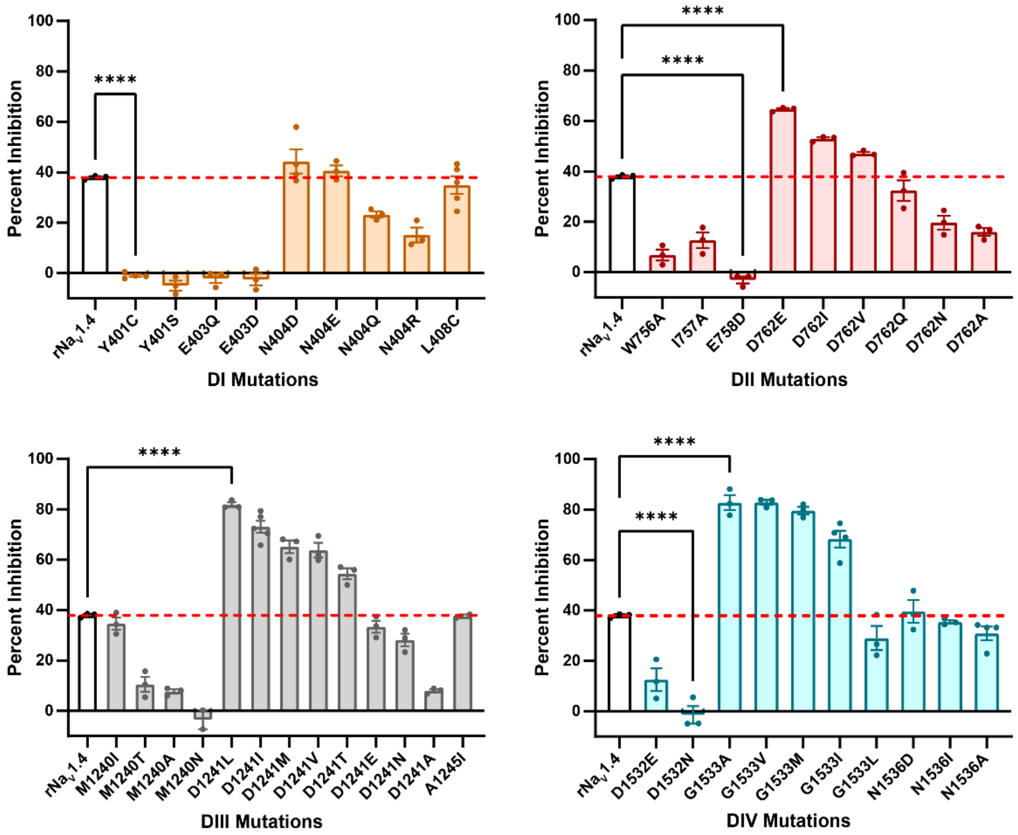


Electrophysiological characterization of 1 µM **2** against select rNa$_V$1.4 DI–DIV mutations. The dashed red line denotes the percent inhibition of WT rNa$_V$1.4 upon application of 1 µM **2** (ca. 38%). Stabilizing mutations show percent inhibition exceeding 38%. Data represent mean ± s.e.m., n ≥ 3. Statistics were calculated with ordinary one-way ANOVA with Dunnett's correction in comparison to WT rNa$_V$1.4. Only **** = p value < 0.0001 is shown.

**Figure S4.** Potency of STX against rNa$_V$1.4 mutant channels.

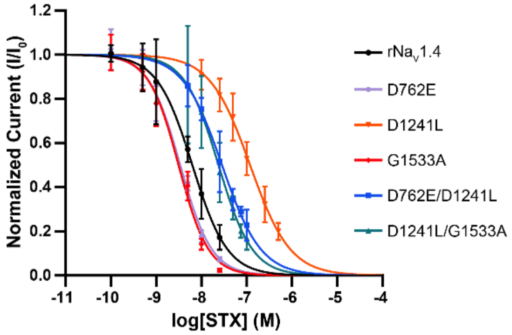


| Channel | $IC_{50}$ | Hill Slope | Fold Change |
|---|---|---|---|
| WT rNa$_V$1.4 | 6.3 ± 0.3 | −1.21 ± 0.07 | - |
| D1241L | 118 ± 6 | −0.98 ± 0.05 | 0.05 |
| G1533A | 3.1 ± 0.2 | −1.31 ± 0.20 | 2 |
| D762E | 4.0 ± 0.2 | −1.25 ± 0.20 | 1.6 |
| D726E/D1241L | 28.4 ± 1.1 | −1.01 ± 0.04 | 0.22 |
| D1241L/G1533A | 23.8 ± 1.4 | −1.16 ± 0.08 | 0.26 |

Concentration-response curves for STX against rNa$_V$1.4 wild-type, D762E, D1241L, G1533A, D762E/D1241L and D1241L/G1533A channels. $IC_{50}$ values ± s.e.m. (nM), Hill coefficients ± s.e.m.. Graph represents mean ± 95% C.I. (for rNa$_V$1.4, n = 3; D762E, n = 4; D1241L, n = 4; G1533A, n = 4; D762E/D1241L, n = 3; D1241L/G1533A, n = 3). The potency of STX towards D1241I was previously reported and is equivalent to that of D1241L.(1)

**Figure S5.** Activation and inactivation parameters for Na$_{V}$1.1–1.7.

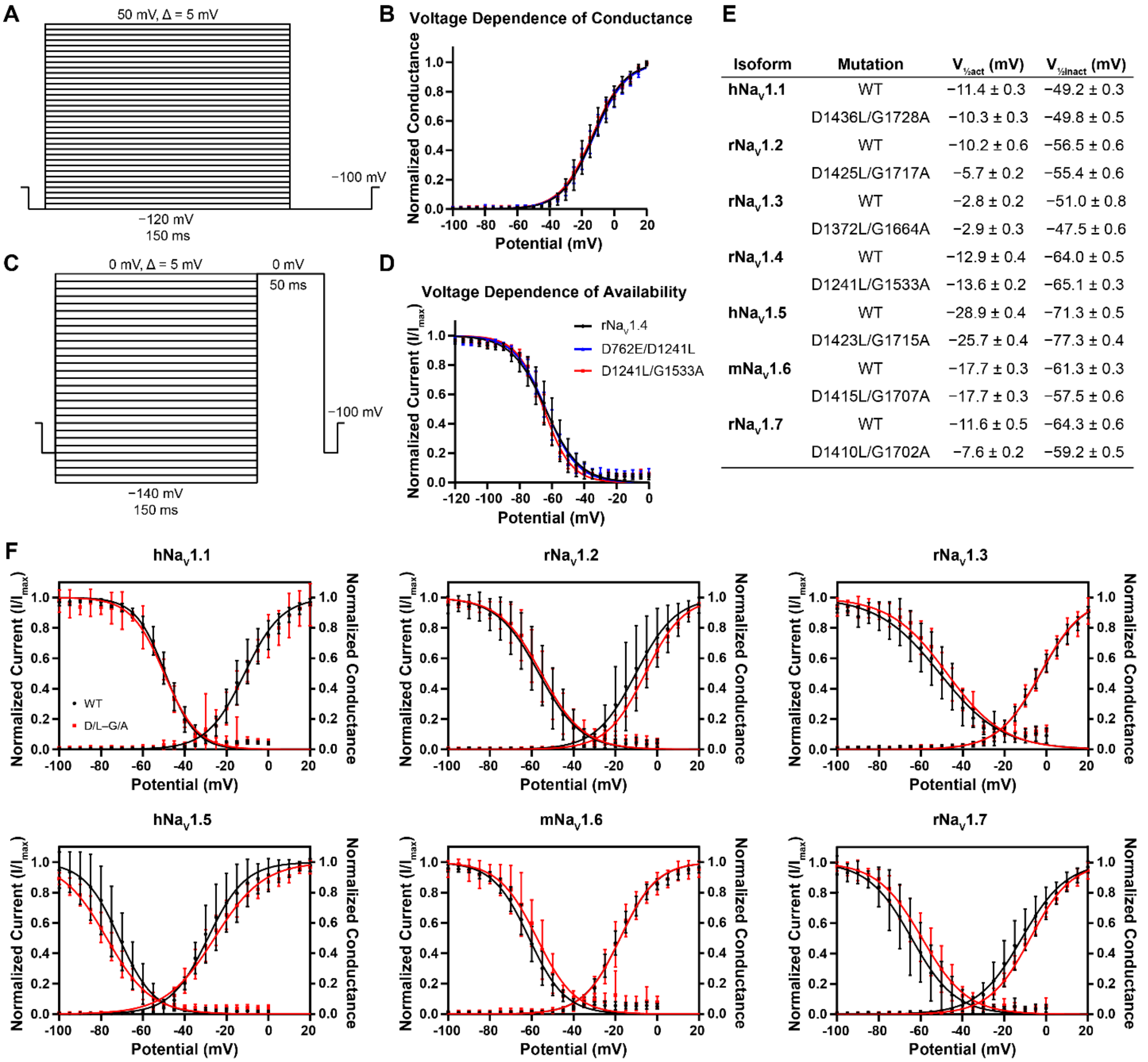


| Isoform | Mutation | $V_{½act}$ (mV) | $V_{½inact}$ (mV) |
|---|---|---|---|
| hNa$_{V}$1.1 | WT | −11.4 ± 0.3 | −49.2 ± 0.3 |
| | D1436L/G1728A | −10.3 ± 0.3 | −49.8 ± 0.5 |
| rNa$_{V}$1.2 | WT | −10.2 ± 0.6 | −56.5 ± 0.6 |
| | D1425L/G1717A | −5.7 ± 0.2 | −55.4 ± 0.6 |
| rNa$_{V}$1.3 | WT | −2.8 ± 0.2 | −51.0 ± 0.8 |
| | D1372L/G1664A | −2.9 ± 0.3 | −47.5 ± 0.6 |
| rNa$_{V}$1.4 | WT | −12.9 ± 0.4 | −64.0 ± 0.5 |
| | D1241L/G1533A | −13.6 ± 0.2 | −65.1 ± 0.3 |
| hNa$_{V}$1.5 | WT | −28.9 ± 0.4 | −71.3 ± 0.5 |
| | D1423L/G1715A | −25.7 ± 0.4 | −77.3 ± 0.4 |
| mNa$_{V}$1.6 | WT | −17.7 ± 0.3 | −61.3 ± 0.3 |
| | D1415L/G1707A | −17.7 ± 0.3 | −57.5 ± 0.6 |
| rNa$_{V}$1.7 | WT | −11.6 ± 0.5 | −64.3 ± 0.6 |
| | D1410L/G1702A | −7.6 ± 0.2 | −59.2 ± 0.5 |

(*A*) Stimulation protocol used to determine the voltage dependence of conductance. Cells were held –100 mV, hyperpolarized to –120 mV for 5 ms, stepped in 5 mV increments from −120 mV to +50 mV (150 ms), and returned to −120 mV (50 ms) before resetting to –100 mV.

(*B*) Voltage dependence of conductance for WT rNa$_{V}$1.4, D762E/D1241L and D1241L/G1533A. $V_{½act}$ ± s.e.m. (mV) or D762E/D1241L = −12.4 ± 0.3. Graphs represent mean ± 95% C.I. (WT rNa$_{V}$1.4, n = 11; rNa$_{V}$1.4 D762E/D1241L, n = 12; rNa$_{V}$1.4 D1241L/G1533A, n = 15).

(*C*) Stimulation protocol used to determine the voltage dependence of availability (steady-state inactivation). Cells were pulsed to 0 mV following a conditioning step between −140 mV to 0 mV (Δ 5 mV per sweep) for 150 ms.

(*D*) Voltage dependence of availability for WT rNa$_{V}$1.4, D762E/D1241L and D1241L/G1533A. $V_{½inact}$ ± s.e.m. (mV) for D762E/D1241L = −63.8 ± 0.4. Graphs represent mean ± 95% C.I. (WT rNa$_{V}$1.4, n = 12; rNa$_{V}$1.4 D762E/D1241L, n = 12; rNa$_{V}$1.4 D1241L/G1533A, n = 14).

(*E*) Table of $V_{½,act}$ and $V_{½,inact}$ for Na$_{V}$1.1–1.7 and the corresponding D/L–G/A double mutant channels. Data represent mean ± s.e.m.

(*F*) Voltage dependence of conductance and voltage dependence of availability for $Na_V$1.1–1.3, 1.5–1.7. Graphs represent mean ± 95% C.I. (h$Na_V$1.1: WT n =6, D/L–G/A n = 3; r$Na_V$1.2: WT n = 5, D/L–G/A n = 9; r$Na_V$1.3: WT n = 6, D/L–G/A n = 6; h$Na_V$1.5: WT n = 3–4, D/L–G/A n = 8–9; m$Na_V$1.6: WT n = 8–9, D/L–G/A n = 4–5; r$Na_V$1.7: WT n = 6–7, D/L–G/A n = 4–5).

**Figure S6.** Ion selectivity of WT rNa$_V$1.4 and rNa$_V$1.4 D762E/D1241L and D1241L/G1533A double mutant channels.

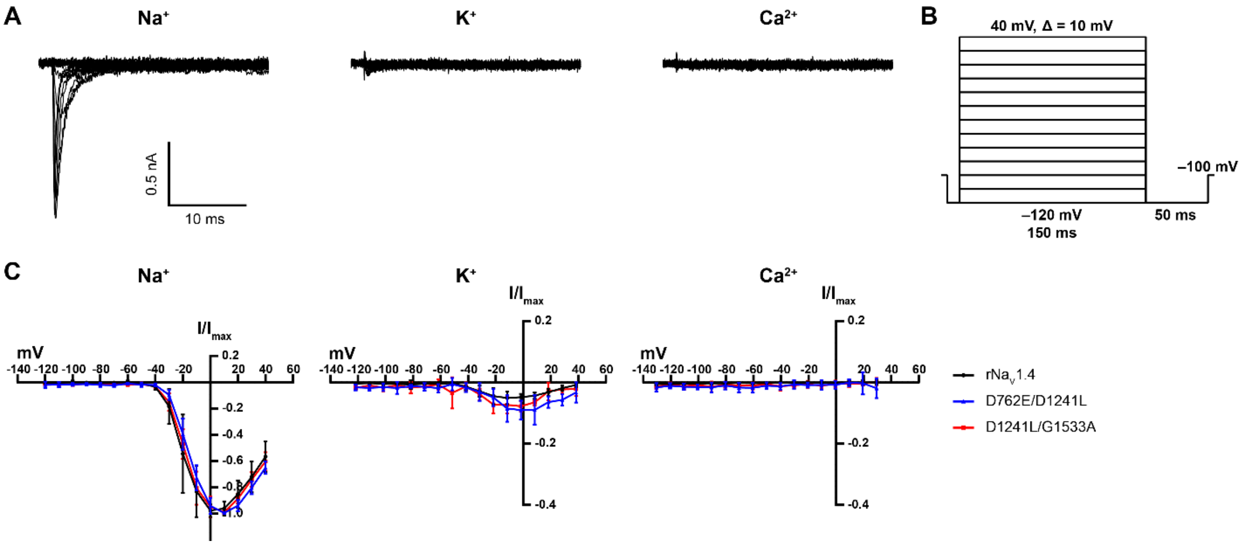


(*A*) Representative current traces for rNa$_V$1.4 D1241L/G1533A transiently expressed in CHO-K1 cells in 160 mM $Na^+$, 160 mM $K^+$, or 110 mM $Ca^{2+}$ external. Each trace is recorded from the same cell.

(*B*) Stimulation protocol for ion selectivity experiments. Cells were held at –100 mV, hyperpolarized to –120 mV (5 ms), stepped in 10 mV increments from −120 mV to +40 mV (150 ms), and returned to −120 mV (50 ms) before resetting to –100 mV.

(*C*) Normalized current-voltage relationships from WT rNa$_V$1.4 and rNa$_V$1.4 D762E/D1241L and D1241L/G1533A in the presence of $Na^+$, $K^+$, and $Ca^{2+}$ external. Each current value is normalized to the peak $Na^+$ current for that cell. There is no recorded $Ca^{2+}$ current at any voltage from the three channels tested. Data represent mean ± 95% C.I. (rNa$_V$1.4, n = 5; rNa$_V$1.4 D762E/D1241L, n = 6; rNa$_V$1.4 D1241L/G1533A, n = 6).

**Figure S7.** Electrophysiological recording data for STX-N$^{m}$Tl **4** against Na$_V$1.1–1.3 and Na$_V$1.5–1.7.

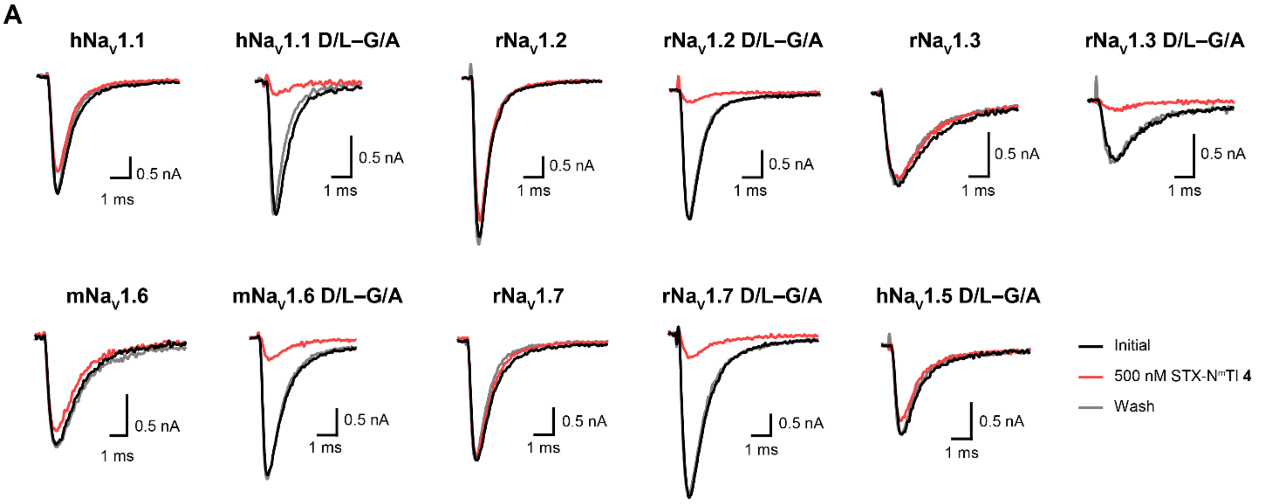


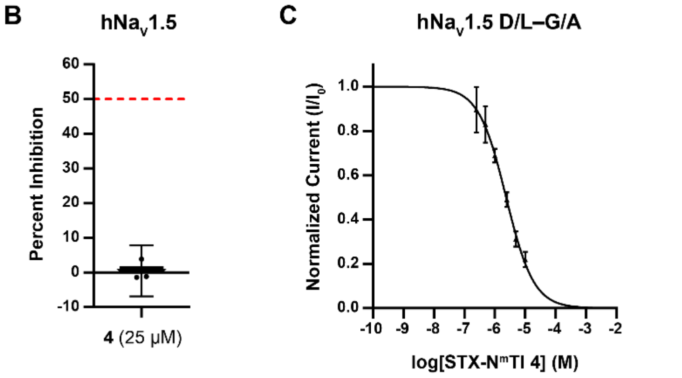


**D**

| Isoform | Mutation | Hill Slope |
|---|---|---|
| **hNa$_V$1.1** | WT | −1.03 ± 0.30 |
| | D1436L/G1728A | −0.82 ± 0.08 |
| **rNa$_V$1.2** | WT | −1.05 ± 0.03 |
| | D1425L/G1717A | −0.84 ± 0.05 |
| **rNa$_V$1.3** | WT | −0.91 ± 0.05 |
| | D1372L/G1664A | −0.83 ± 0.05 |
| **hNa$_V$1.5** | WT | nd |
| | D1423L/G1715A | −0.96 ± 0.04 |
| **mNa$_V$1.6** | WT | −1.02 ± 0.03 |
| | D1415L/G1707A | −0.90 ± 0.07 |
| **rNa$_V$1.7** | WT | −1.23 ± 0.10 |
| | D1410L/G1702A | −0.81 ± 0.04 |

(*A*) Representative voltage-clamp traces for WT and mutant Na$_V$1.1–1.3, 1.5–1.7 with 500 nM of STX-N$^{m}$Tl **4** applied. Traces collected in the order: Initial, 500 nM **4**, Wash.
(*B*) Percent inhibition of WT hNa$_V$1.5 by 25 µM of STX-N$^{m}$Tl **4**. Data represents mean ± 95% C.I. (n = 3).
(*C*) Concentration-response curve for **4** with hNa$_V$1.5 D1423L/G1715A. Data represents mean ± 95% C.I. (n = 4).
(*D*) Table summary of WT Na$_V$1.1–1.7 isoforms, the corresponding double mutant channels, and Hill slopes of concentration-response curves. Hill slopes ± s.e.m.; nd = not determined.

### SI Appendix, Materials and Methods

#### Synthesis

All reagents were obtained commercially. Organic solutions were concentrated under reduced pressure (ca. 60 Torr) by rotary evaporation. HPLC-grade $CH_3CN$ was obtained from commercial suppliers and used as is. *N,N*-Dimethylformamide (DMF) was passed through two columns of activated alumina prior to use.

Semi-preparative high-performance liquid chromatography (HPLC) was performed on a Varian ProStar model 210. High-resolution mass spectra were obtained from the Vincent Coates Foundation Mass Spectrometry Laboratory at Stanford University. Samples were analyzed by LC-flow injection ESI/MS on the Waters Acquity H-Class Plus UPLC and Thermo Exploris 240 BioPharma Orbitrap mass spectrometer scanning *m/z* 100–1000 Da. A carrier solvent of methanol or acetonitrile with 0.1% formic acid at a flow rate of 0.2mL/min was used to transport the injected sample to the source directly, without chromatographic separation.

Saxitoxin derivatives were quantified by $^1H$ NMR spectroscopy on a Varian Inova 600 MHz NMR instrument using distilled DMF as an internal standard. A relaxation delay ($d_1$) of 20 s and an acquisition time (at) of 10 s were used for spectral acquisition. The concentration of the toxin derivative was determined by integrating $^1H$ signals corresponding to the toxin and a fixed concentration of the DMF standard.

STX-amides can be obtained through modification of a previously described synthesis (*J. Am. Chem. Soc.* **2008**, *130*, 12630–12631).(2) The desired ligands were purified by reversed-phase HPLC (0–60% $CH_3CN$ in 10 mM aqueous trifluoroacetic acid over 60 min unless otherwise noted). Fractions were collected, frozen, and lyophilized to remove all volatiles. Fractions were quantified by $^1H$ NMR integration and lyophilized again to yield the desired bis-guanidinium derivative.

**STX-NAc 1**

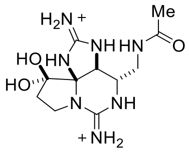

At a flow rate of 4 mL/min and gradient 0–10% $CH_3CN$ in $H_2O$/trifluoroacetic acid over 20 min, STX-NAc **1** had retention time of 4.91 min and was isolated as a white powder following lyophilization (0.71 µmol, 48%, $^1$H NMR quantitation).

**$^1$H NMR** (600 MHz, $D_2O$) δ 4.73 (s, 1H), 3.83 (t, $J$ = 10.0 Hz, 1H), 3.71–3.66 (m, 1H), 3.66–3.58 (m, 2H), 3.23 (dd, $J$ = 14.0, 4.6 Hz, 1H), 2.47 (dd, $J$ = 14.1, 8.0 Hz, 1H), 2.40–2.33 (m, 1H), 2.03 (s, 3H) ppm

**HRMS** ($ESI^+$) calcd for $C_{11}H_{19}N_7O_3$ 297.1549 found 298.1621 ($MH^+$)

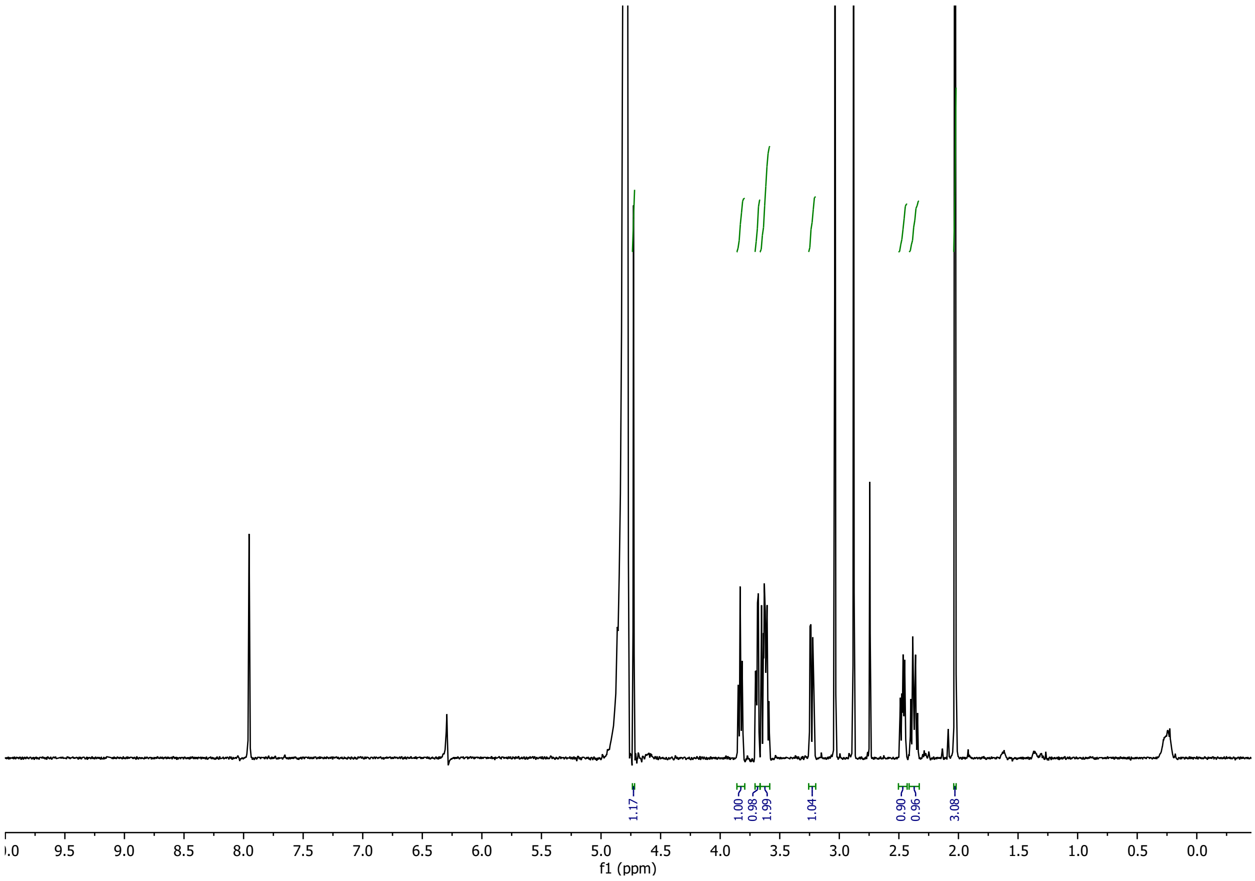

**STX-NBz 2**

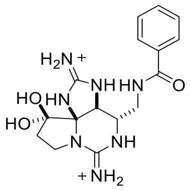


At a flow rate of 4 mL/min, STX-NBz **2** had retention time of 18.61 min and was isolated as a white powder following lyophilization (0.57 µmol, 36%, $^{1}$H NMR quantitation)

**$^{1}$H NMR** (600 MHz, $D_2O$) δ 7.79 (d, $J$ = 7.7 Hz, 2H), 7.67 (t, $J$ = 7.5 Hz, 1H), 7.57 (dd, $J$ = 7.7, 7.5 Hz, 2H), 3.88–3.81 (m, 2H), 3.78 (t, $J$ = 9.9 Hz, 1H), 3.56 (dt, $J$ = 9.3, 8.8 Hz, 1H), 3.40 (dt, $J$ = 9.3, 8.4 Hz, 1H), 2.46–2.40 (m, 1H), 2.39 – 2.32 (m, 1H) ppm

*H5 obscured by HDO peak ((its present though))

**HRMS** (ESI$^{+}$) calcd for $C_{16}H_{21}N_7O_3$ 359.1706 found 360.1779 ($MH^{+}$)

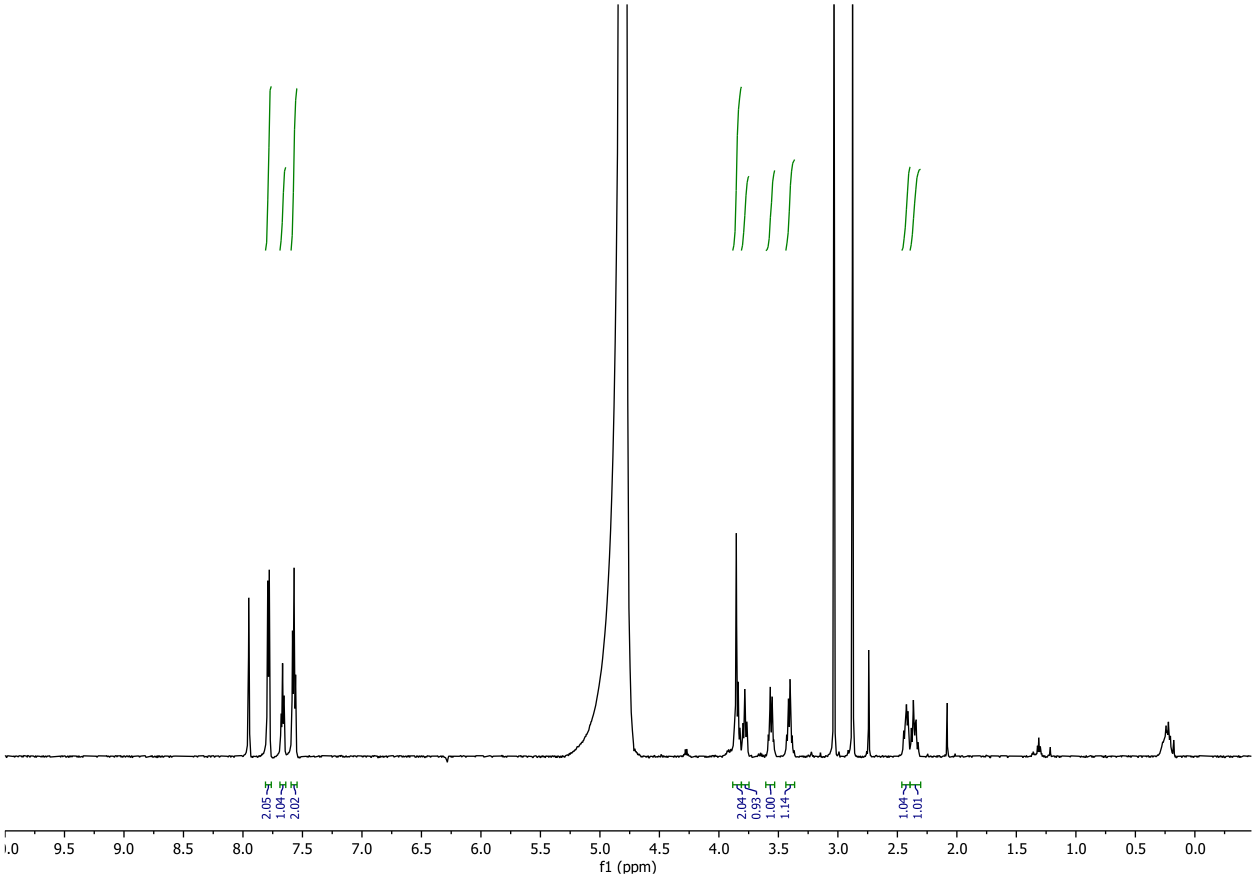

**STX-N$^{p}$Tl 3**

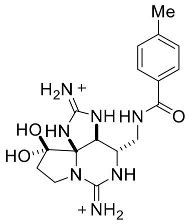


At a flow rate of 4 mL/min, STX-N$^{p}$Tl **3** had retention time of 17.51 min and was isolated as a white powder following lyophilization (0.21 μmol, 13%, $^{1}$H NMR quantitation)

**$^{1}$H NMR** (600 MHz, $D_2O$) δ 7.64 (d, *J* = 8.0 Hz, 2H), 7.35 (d, *J* = 8.0 Hz, 2H), 3.82–3.70 (m, 3H), 3.51 (dd, *J* = 12.9, 3.3 Hz, 1H), 3.34 (dt, *J* = 9.3, 9.3 Hz, 1H), 2.40–2.35 (m, 4H), 2.34–2.27 (m, 1H) ppm

*H5 obscured by HDO peak

**HRMS** (ESI$^{+}$) calcd for $C_{17}H_{23}N_7O_3$ 373.1862 found 374.1942 ($MH^{+}$)

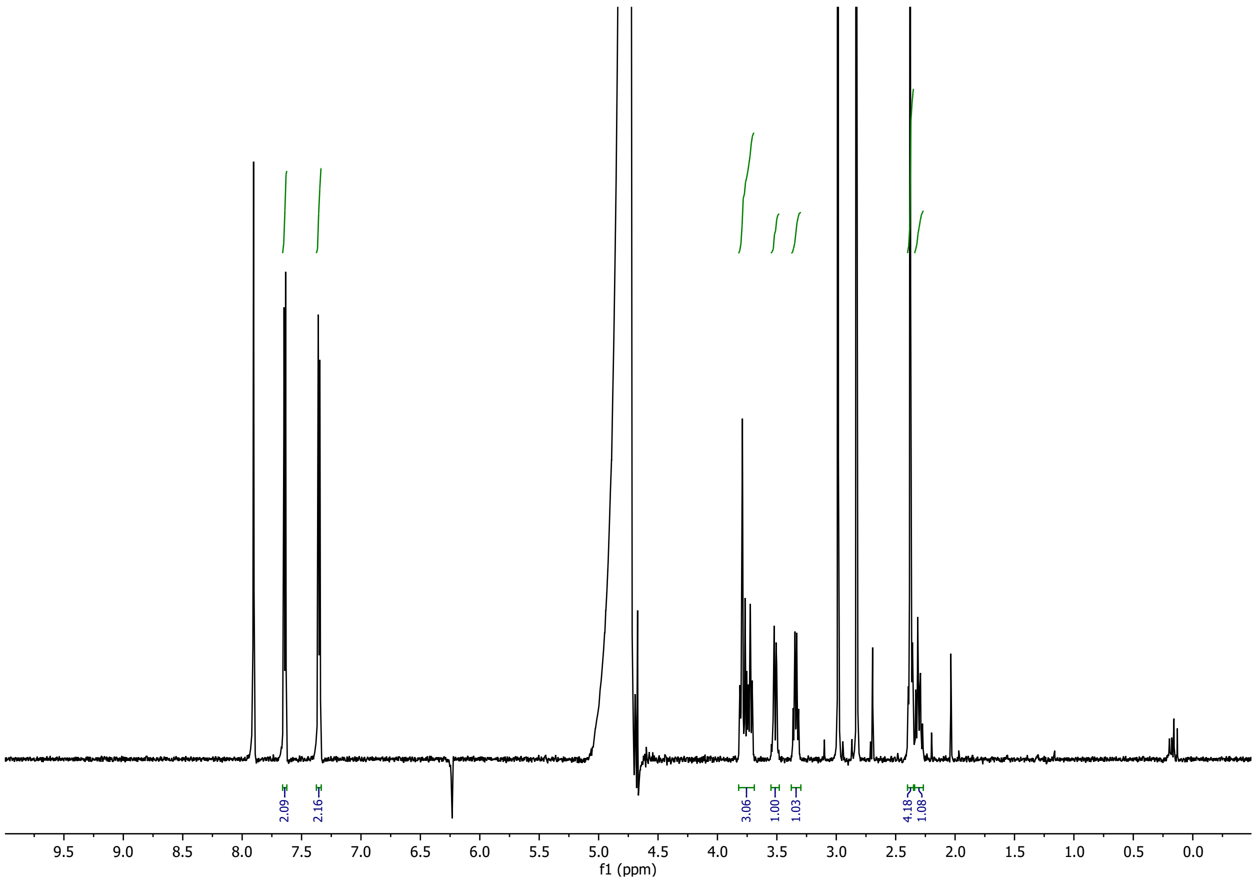

**STX-N$^{m}$Tl 4**

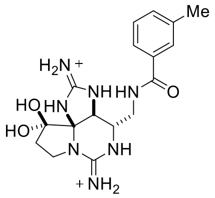


At a flow rate of 4 mL/min, STX-N$^{m}$Tl **4** had retention time of 17.56 min and was isolated as a white powder following lyophilization (0.20 µmol, 14%, $^{1}$H NMR quantitation)

**$^{1}$H NMR** (600 MHz, $D_2O$) δ 7.62 (s, 1H), 7.58 (d, $J$ = 7.6 Hz, 1H), 7.51 (d, $J$ = 7.6 Hz, 1H), 7.46 (t, $J$ = 7.6 Hz, 1H), 4.84 (s, 1H), 3.87–3.81 (m, 2H), 3.78 (t, $J$ = 10.1 Hz, 1H), 3.60–3.52 (m, 1H), 3.41 (dt, $J$ = 9.9, 8.1 Hz, 1H), 2.46–2.40 (m, 4H), 2.36 (dt, $J$ = 13.9, 9.9 Hz, 1H) ppm

**HRMS** (ESI$^{+}$) calcd for $C_{17}H_{23}N_7O_3$ 373.1862 found 374.1942 ($MH^{+}$)

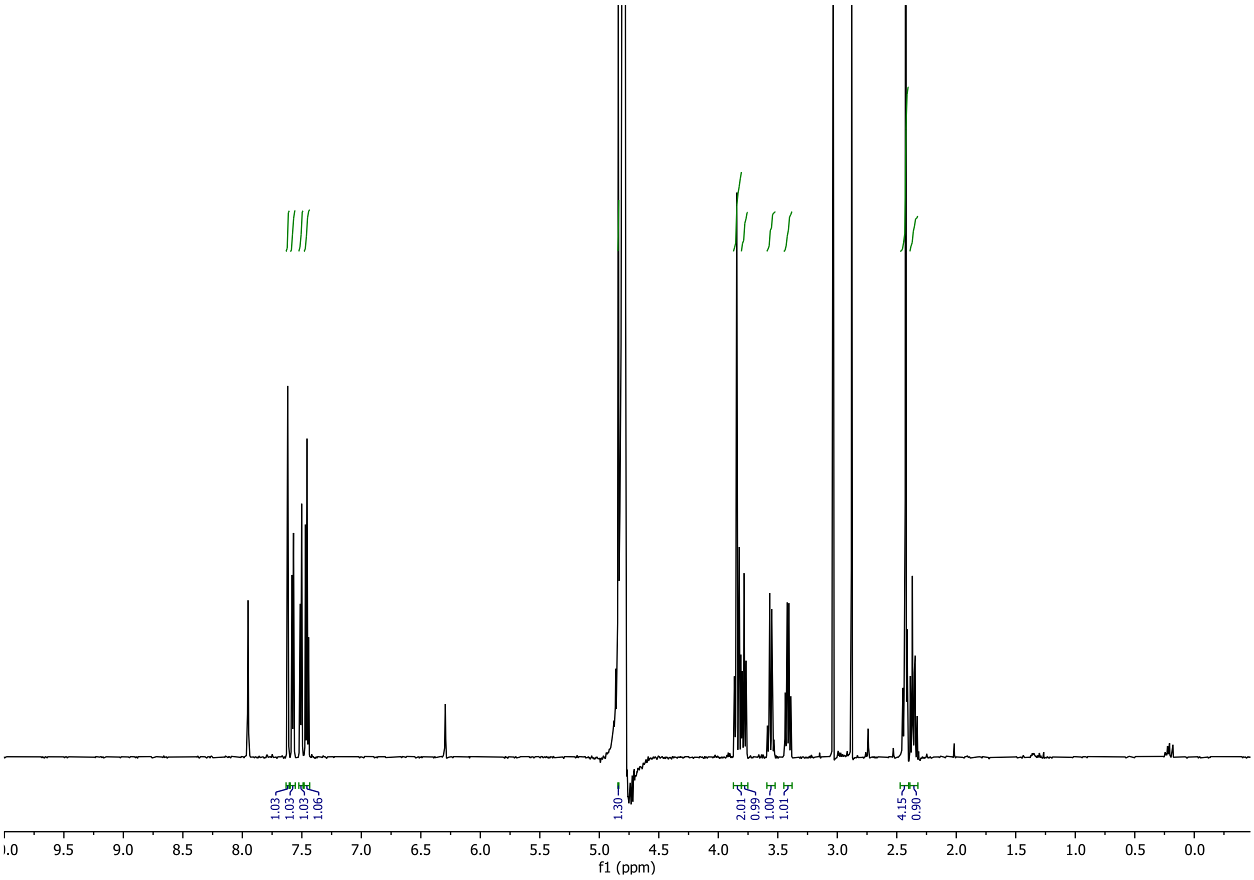

**STX-M$^{o}$Tl 5 ***

*This compound was synthesized through an alternative route based on *J. Am. Chem. Soc.* **2008**, *130*, 12630–12631.(2)

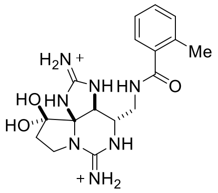


At a flow rate of 4 mL/min and gradient 0–50% $CH_3CN$ in $H_2O$/heptafluorobutyric acid over 50 min, STX-M$^{o}$Tl **5** had retention time of 35.49 min.

**$^1$H NMR** (600 MHz, $D_2O$) δ 7.47 (dd, $J$ = 7.7, 7.6 Hz, 1H), 7.43 (d, $J$ = 7.7 Hz, 1H), 7.37 (d, $J$ = 7.7 Hz, 1H), 7.34 (dd, $J$ = 7.7, 7.6 Hz, 1H), 4.84 (s, 1H), 3.88–3.79 (m, 3H), 3.56–3.46 (m, 2H), 2.47–2.41 (m, 1H), 2.41–2.33 (m, 4H) ppm

**HRMS** ($ESI^+$) calcd for $C_{17}H_{23}N_7O_3$ 373.1862 found 374.1934 ($MH^+$)

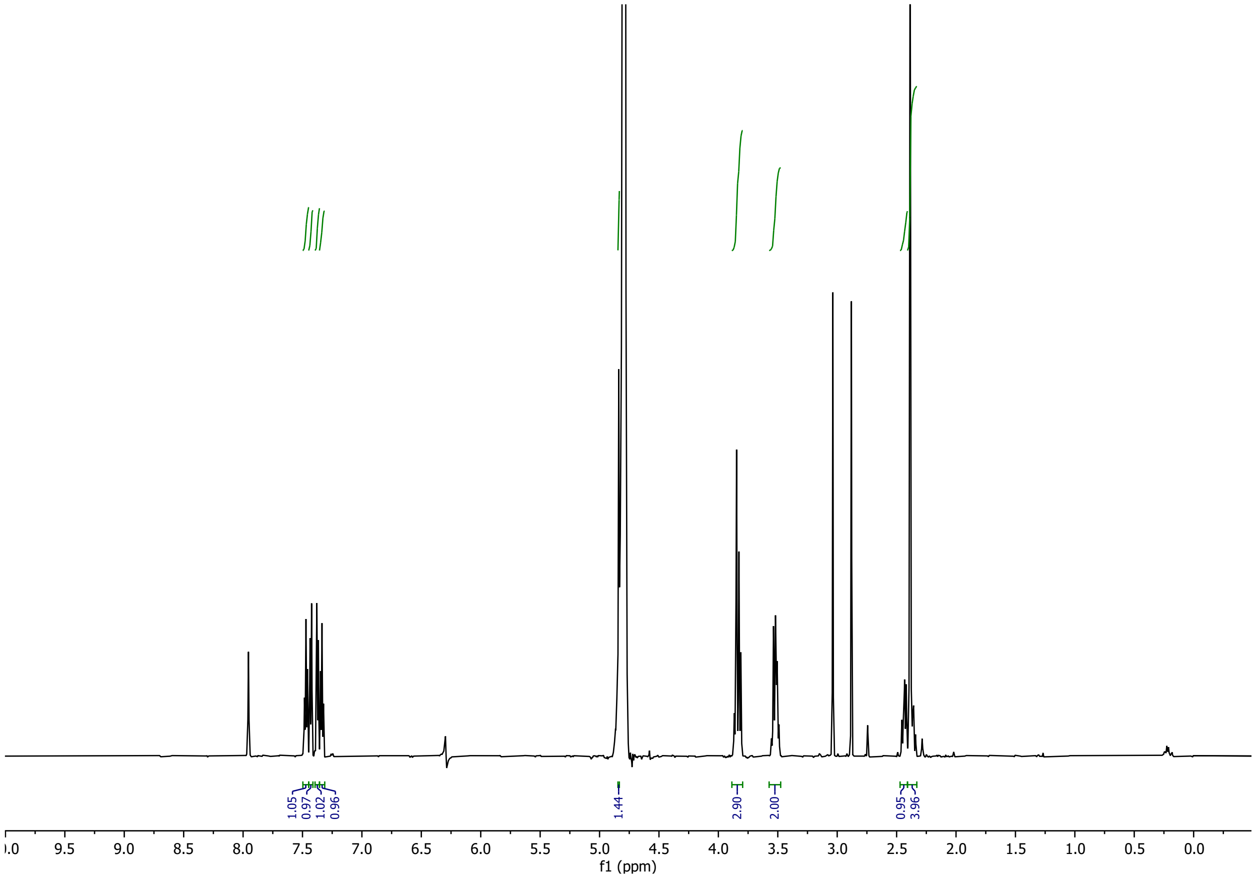

**STX-NCy 6**

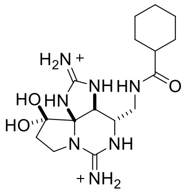


At a flow rate of 4 mL/min, STX-NCy **6** had retention time of 16.60 min and was isolated as a white powder following lyophilization (0.56 µmol, 36%, $^1$H NMR quantitation)

**$^1$H NMR** (600 MHz, $D_2O$) δ 4.73 (s, 1H), 3.84 (dt, $J$ = 10.2, 1.8 Hz, 1H), 3.7–3.64 (m, 2H), 3.63–3.57 (m, 1H), 3.23 (dt, $J$ = 9.9, 9.2 Hz, 1H), 2.50–2.44 (m, 1H), 2.37 (dt, $J$ = 14.0, 9.9 Hz, 1H), 2.29–2.21 (m, 1H), 1.84–1.75 (m, 4H), 1.68 (d, $J$ = 12.8 Hz, 1H), 1.40–1.25 (m, 4H), 1.25–1.16 (m, 1H) ppm

**HRMS** (ESI$^+$) calcd for $C_{16}H_{27}N_7O_3$ 365.2175 found 366.2246 ($MH^+$)

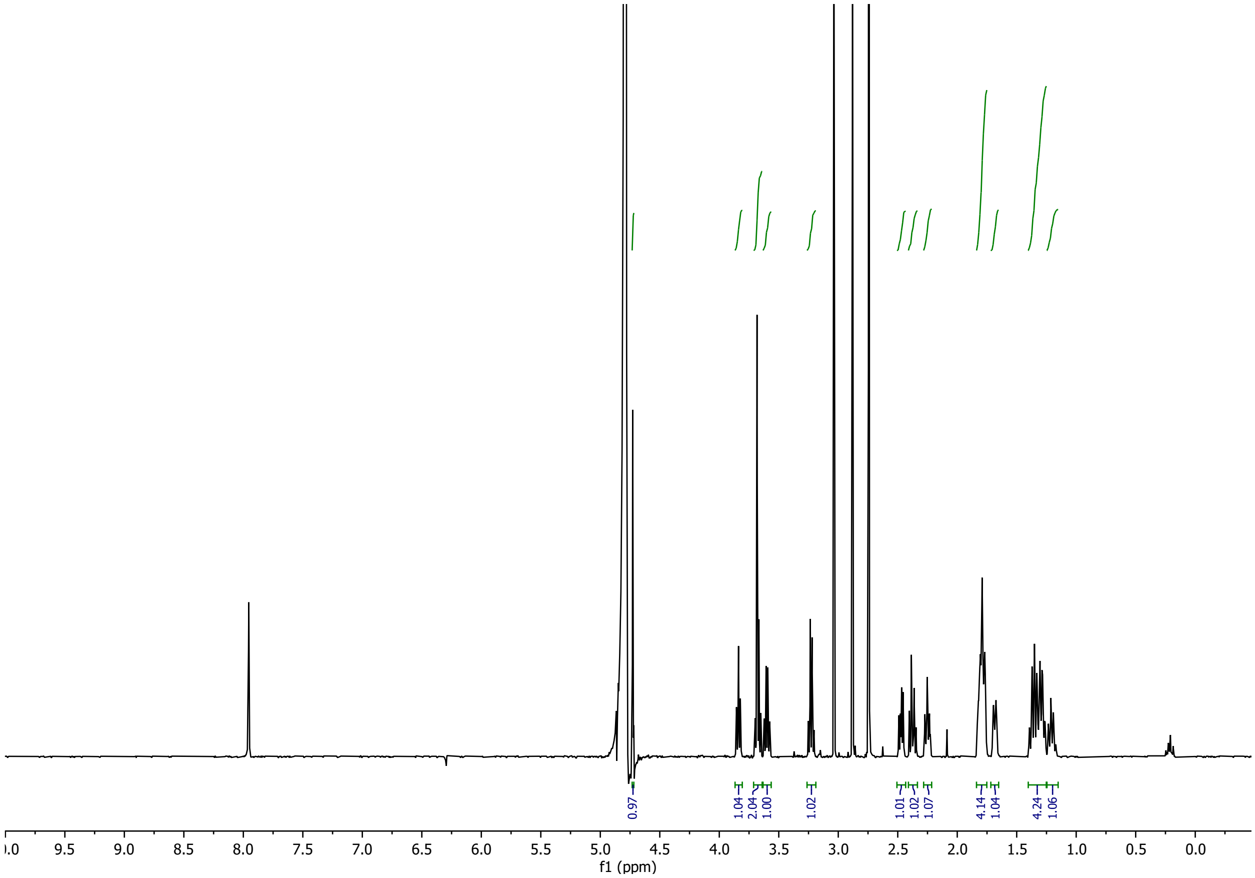

**Mutagenesis**
Several rNa$_V$1.4 mutants were previously reported:

- Y401S, E403D, W756A, I757A, E758D, M1240I, M1240A, M1240N, D1241T, D1241E, D1241N, D1241A, D1532E(1, 3)
- M1240T, D1241I(4)
- D762E, G1533A, E403Q(5)
- D1532N(6, 7)
- Y401C(7, 8)

Forward primer sequences 5' to 3':
rNa$_V$1.4:
Domain I

- E403Q: CTCATGACGCAGGACTACTGGCAGAACCTTTTCCAGC
- N404D: GCAGGACTACTGGGAGGACCTTTTCCAGCTGACC
- N404E: GCAGGACTACTGGGAGGAGCTTTTCCAGCTGACC
- N404Q: GCAGGACTACTGGGAGCAGCTTTTCCAGCTGACC
- N404R: GCAGGACTACTGGGAGCGCCTTTTCCAGCTGACC
- L408C: GGAGAACCTTTTCCAGTGCACCCTACGAGCTGCTG

Domain II

- D762E: AATGGATCGAGACCATGTGGGAGTGCATGGAGGTGG
- D762V: GATCGAGACCATGTGGGTCTGCATGGAGGTGGCC
- D762I: GGATCGAGACCATGTGGATCTGCATGGAGGTGGCC
- D762Q: GGATCGAGACCATGTGGCAGTGCATGGAGGTGG
- D762N: GGATCGAGACCATGTGGAACTGCATGGAGGTGGC
- D762A: GATCGAGACCATGTGGGCCTGCATGGAGGTGGC

Domain III

- D1241L: CATTCAAGGGTTGGATGCTGATCATGTATGCAGCTGTGG
- D1241M: CCACATTCAAGGGTTGGATGATGATCATGTATGCAGCTGTGG
- D1241V: CCACATTCAAGGGTTGGATGGTGATCATGTATGCAGCTGTGG
- A1245I: GGGTTGGATGGATATCATGTATATCGCTGTGGACTCCCGG

Domain IV

- G1533A: CCGGCTGGGACGCTCTTCTGAACCCCATCC
- G1533V: CCGGCTGGGACGTGCTTCTGAACCCCATCC
- G1533M: CCGGCTGGGACATGCTTCTGAACCCCATCCTCAAC
- G1533I: CCGGCTGGGACATTCTTCTGAACCCCATCCTCAAC
- G1533L: CCGGCTGGGACCTGCTTCTGAACCCCATCCTCAAC
- N1536D: CTGGGACGGGCTTCTGGACCCCATCCTCAACAG

- N1536I: GCTGGGACGGGCTTCTGATCCCCATCCTCAACAG
- N1536A: TGGGACGGGCTTCTGGCCCCCATCCTCAACAG

hNa$_V$1.1:

- D1436L: GCCACATTCAAAGGATGGATGCTCATAATGTATGCAGCAG
- G1728A: CCTGCTGGCTGGGATGCTTTGCTAGCACCCATTCTC

rNa$_V$1.2:

- G1717A: CTGCGGGCTGGGACGCTCTGCTGGCCCCT
- D1425L: GCCACCTTTAAAGGATGGATGCTGATCATGTATGCAGCTGTTG

hNa$_V$1.5:

- D1423L: CAACATTTAAAGGCTGGATGCTCATTATGTATGCAGCTGTCG
- G1715A: GGCCGGCTGGGATGCTCTCCTCAGCCCCAT

mNa$_V$1.6:

- D1415L: CCTTCAAAGGCTGGATGCTCATCATGTATGCAGCTGTAG
- G1707A: CCTCTGCTGGTTGGGATGCTTTACTGCTGCCAATCC

**Sodium Current Recordings**

*Kinetic Measurements of Block*

Currents were elicited by 10 ms step depolarizations from –100 mV to 0 mV at a rate of 0.5 Hz. Using a pinch valve perfusion system (Automate Scientific, Berkeley, CA) to control the perfusion rate at 1.5 mL/min, 1.5 mL of STX or STX-N$^{m}$Tl **4** was washed onto cells and equilibrated for 1 min. Cells were then washed with 1–2 mL of external solution until full return of current. Only cells that returned within 10% of initial peak current were used in the analysis. A ligand concentration that approximated the respective $IC_{50}$ value was used to measure the rate of binding (5 nM STX in WT, 4.5 µM **4** in WT, and 25 nM **4** in rNa$_V$1.4 D1241L/G1533A). Peak current was normalized to the initial current and plotted against time. The decay in current during wash on and recovery of current during wash out were fit to exponential curves to obtain the on-rate time constant ($\tau_{on}$) and off-rate time constant ($\tau_{off}$). Rate constants for block ($k_{on}$) and recovery ($k_{off}$) were derived according to the equations by Hahn and Strichartz.(9)

*State-dependence of Binding*

Currents were elicited by a 10 ms test pulse to 0 mV at a frequency of 0.5 Hz. Test pulses were preceded by 200 ms conditioning pulses to –70 mV or –120 mV to sample different conformational states. A baseline involving 5 scans was recorded prior to toxin exposure. Using a perfusion system (Automate Scientific, Berkeley, CA), 1.5 mL of STX or STX-N$^{m}$Tl **4** was washed onto cells at a rate of 1.5 mL/min and allowed to equilibrate for 1 min. Cells were then washed with 1–2 mL of external solution. Only cells that returned within 10% of initial peak current were used for analysis. The baseline and ligand-treated peak current were recorded as the average of 5 scans.

*Voltage-dependence of Conductance (Activation)*

Currents were evoked using the stimulus protocol depicted in Figure S5A at sampling rate of 50 kHz. Peak current was converted to conductance (G) using the equation $I_{Na}/(V_m–V_{rev})$, where $I_{Na}$ is the measured peak sodium current, $V_m$ is the applied membrane potential, and $V_{rev}$ is the experimentally determined reversal potential for sodium. Conductance at each potential step was normalized to the maximum conductance and plotted against voltage. The G–V curve was fit to a sigmoidal curve constrained to [0,1] with variable slope and the midpoint of the curve, $V_{1/2act}$, was calculated in GraphPad Prism. Cells with >0.5 nA current and experimentally determined $V_{rev}$ between 32–38 mV, within 3 mV of the calculated $V_{rev}$, were used for analysis.

*Voltage-dependence of Availability (Steady-state Inactivation)*

Currents were evoked using the stimulus protocol depicted in Figure S5C at a sampling rate of 100 kHz. Peak current was normalized to the largest current across scans and plotted against voltage. The I–V curve was fit to a sigmoidal curve constrained to [0,1] with variable slope and the midpoint of the curve, $V_{1/2inact}$, was calculated in GraphPad Prism. Cells with >0.5 nA current were used for analysis.

*Ion Selectivity*

The internal solution was composed of 40 mM CsF, 1 mM EGTA, 20 mM HEPES, and 125 mM CsCl (pH 7.4 with 50 wt. % aqueous CsOH, 325 mOsm); the external solution comprised 160 mM X-Cl (X = Na, K) or 110 mM $XCl_2$ (X = Ca), 2 mM $CaCl_2$, and 20 mM HEPES (pH 7.4 with 50 wt. % aqueous CsOH, 325–330 mOsm). Liquid junction potentials (LJP) were zeroed at the start of each experiment in $Na^+$ external. LJPs were calculated and experimental confirmed to be –1.6 mV for $K^+$ and –10.5 mV for $Ca^{2+}$. The LJP was corrected during data analysis.

Cells were first patched in $Na^+$ external solution and equilibrated for at least 5 min after establishing the whole-cell and voltage-clamp configuration. Currents were evoked using the protocol depicted in Figure S6B at a sampling rate of 50 kHz. External solution was replaced through perfusion to either $K^+$ or $Ca^{2+}$ external solution, and currents were evoked by the same protocol.

To minimize leak currents, whole-cell currents were measured as the difference between peak currents in a 13.5–20 ms window and average sustained current was measured between 150–160 ms. Data were filtered during analysis by a 2 kHz low-pass Bessel filter applied in Clampfit. The peak current at each voltage step was normalized to the largest peak current in $Na^+$ external conditions.
